# Pore-size dynamics control complex volume swelling in pyroptosis

**DOI:** 10.1101/2025.03.27.645729

**Authors:** Estelle Bastien, Guillaume Duprez, Hélène Delanoë-Ayari, Hubert Leloup, Charlotte Rivière, Virginie Petrilli, Pierre Recho, Sylvain Monnier

**Author notes:** To whom correspondence should be addressed. E.mail. Conceptualization: E.B, S.M. and P.R.; methodology: E.B., S.M.and P.R; investigation: E.B, G.D. and H.D; writing – original draft: E.B, S.M. and P.R.; draft editing: E.B, S.M., P.R., G.D, C.R., V.P.; funding acquisition: S.M., P.R.,V.P. and C.R.; Supervision: E.B, S.M. and P.R. The authors declare no competing interest. P.R. and S.M. contributed equally to this work.

## Abstract

Pyroptosis, an inflammatory form of cell death, is characterized by massive cell swelling and plasma membrane rupture. Although swelling was recently shown to occur in two steps, the molecular and biophysical mechanisms driving this process remained unclear. Using fast quantitative microscopy, we reveal that between the two swelling phases, cell volume transiently stabilizes despite sustained plasma membrane permeability to ions and small molecules. From a biophysical perspective, the existence of such a plateau is puzzling, as ion pumps should not be able to regulate cell volume under these conditions. To address this, we developed a physical model based on an ion pump and leak framework that incorporates the dynamics of non-selective pore formation. Experimentally, we demonstrate that the plateau phase is controlled by the dynamics of the GSDMD pore enlargement, which is modulated by Ninj1 activation, possibly through intracellular calcium. Ninj1-mediated lesions are also required for the second swelling phase. We further show that fully opened GSDMD pores display an effective hydrodynamic radius slightly above 1.9 nm, providing an *insitu* upper bound for pore size. Together, our findings demonstrate that pyroptotic volume dysregulation emerges from the successive and interdependent actions of GSDMD and Ninj1, each imparting distinct permeability regimes associated with increased water filtration and decreased ion selectivity due to pore opening. These insights bridge molecular and biophysical perspectives on lytic cell death and may inform the broader understanding of membrane rupture in inflammatory and pathological contexts.

**Significance Statement:** Among programmed modes of lytic cell death, pyroptosis mediated by gasdermin D (GSDMD) and ninjurin-1 (Ninj1) involves dramatic changes in cell shape and large volume fluctuations, fundamentally altering the cell’s physical properties. By combining optogenetics, quantitative microscopy, and modeling, we show that a progressive increase in plasma membrane pore size drives cell swelling and membrane lysis through successive and interdependent actions of GSDMD and Ninj1, each imparting distinct permeability regimes associated with increased water filtration and decreased ion selectivity. A deeper understanding of these dynamic cell modifications will shed light on the molecular and biophysical mechanisms driving different forms of cell death.

---

Cell death, a fundamental biological process, is often accompanied by significant morphological changes. Notably, pyroptosis – a key pro-inflammatory form of cell death – is characterized by drastic cell volume dysregulation with distinct events, including swelling, plasma membrane (PM) blebbing and rupture, ultimately releasing intracellular content that enhances inflammation (1). Pyroptotic swelling arises from ion fluxes and osmotic imbalance due to PM permeabilization mediated by two molecular actors: gasdermin D (GSDMD), which forms pores (2), and the PM protein ninjurin-1 (Ninj1), which generates larger lesions (3). This makes pyroptosis an ideal model to investigate how the dynamic evolution of PM permeability drives cell volume dysregulation.

In canonical pyroptosis, pathogens or danger signals activate innate immune receptors, leading to the assembly of multiprotein complexes known as inflammasomes and the formation of micrometer-scale specks (4). Within these complexes, the adaptor ASC recruits and activates caspase-1 (CASP1), which drives both the maturation of pro-inflammatory cytokines such as IL-1*β* and the cleavage of GSDMD. The resulting N-terminal fragments oligomerize into PM pores (5–7). GSDMD pores release small molecules, including mature IL-1*β*, but do not account for the escape of larger cytosolic components. This step relies on Ninj1-dependent lesions, which amplify PM permeabilization and ultimately drive its rupture (3, 8). To directly probe pore formation while bypassing variability from external cues and upstream signalling, we used our recently developed optogenetic system (opto-ASC) (9) (Fig. 1A). This tool enables precise light-controlled activation of inflammasomes, triggering GSDMD pores and Ninj1 lesions, and faithfully recapitulating the hallmarks of pyroptosis-induced cell swelling.

**Fig. 1.**
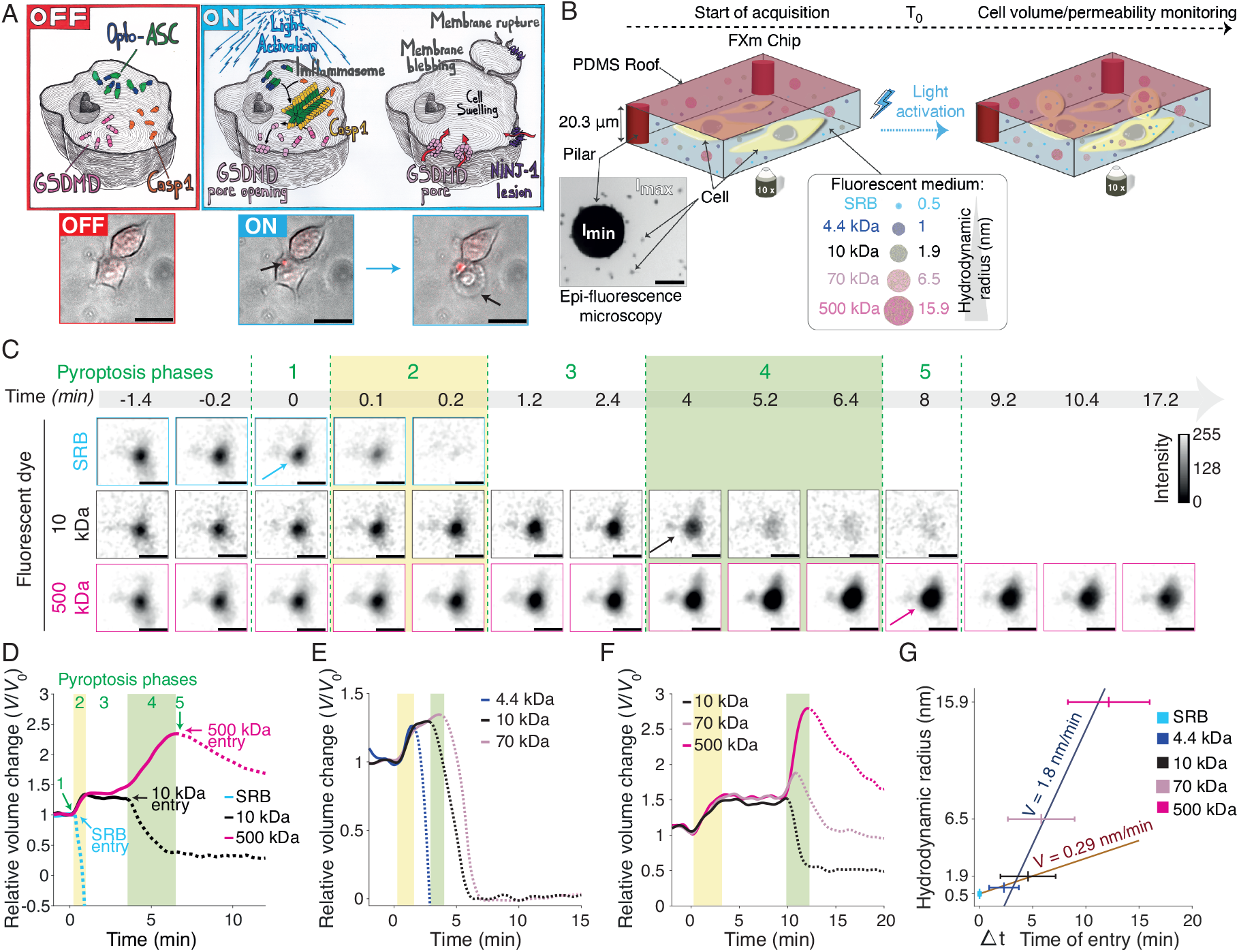
Single-cell Fluorescence eXclusion microscopy analysis of pyroptosis uncovers swelling dynamics and pore size expansion. (A) Schematic of the optogenetic iBMDM model for controlled pyroptosis induction. Without blue light (OFF), cells express GSDMD, inactive Casp-1 (orange), and the optogenetic construct opto-ASC (ASC–CRY2–TagRFP). Blue light activation (ON) triggers CRY2-dependent ASC oligomerization, recruitment of activated Casp-1 (yellow), inflammasome speck formation, and GSDMD cleavage. The GSDMD N-terminal fragments insert into the plasma membrane (PM), forming pores that drive cell swelling, PM blebbing and rupture via Ninj1 (9). Representative merged images show the successive inflammasome specks (red puncta corresponding to the RFP signal of ASC–CRY2–TagRFP) and PM blebbing formation (arrow). (B) Single-cell volume measurement by Fluorescence eXclusion microscopy (FXm). Cells were seeded in a PDMS chamber containing fluorescent probes that cannot freely enter the cytoplasm. The chamber includes micropillars that define its height and provide the zero-fluorescence reference (*I*_min_). As a result, cells appear as darker regions compared with the surrounding medium (*I*_max_) in epi-fluorescence images (10× objective, arrow). Because fluorescence intensity is linearly proportional to the excluded volume height, integrating the difference between the background and local intensities over the projected cell area yields an absolute measurement of cell volume as long as the probe cannot permeate the PM (35) (see Materials & Methods and Sup Fig. 1). After optogenetic activation by blue light illumination, excluded volumes were tracked over time using fast time-lapse acquisitions (8 s). To monitor pore opening, the chamber was filled with fluorescent probes of increasing hydrodynamic radii (Sulforhodamine B (SRB), 0.5 nm, to 500 kDa Dextran, 15.9 nm (36, 37)). Probe entry of increasing size revealed the minimal pore size at the PM, detected as an increase in intracellular fluorescence together with a corresponding decrease in the excluded cell volume. (C) Examples of FXm timelapse images (scale bar, 20 *µ*m; arrows indicate probe entry into cells). (D) Representative normalized dynamics of apparent cell volume during pyroptosis in the presence of SRB (blue), 10 kDa Dextran-AF647 (black), and 500 kDa Dextran-FITC (pink), relative to the initial volume *V*_0_ at the onset of swelling (*t*_0_). Arrows indicate probe entry, detected as a drop in the excluded cell volume curve measured with the corresponding probe. Five distinct phases are indicated along the apparent cell volume curves: (1) onset of swelling linked to the first PM permeabilization marked by SRB entry; (2) early swelling (yellow); (3) volume plateau; (4) late swelling (green), marked by 10 kDa dextran entry, and (5) PM rupture, marked by 500 kDa Dextran entry, defining *V*_max_. (E-F) Representative normalized dynamics of apparent cell volume in the presence of two sets of fluorescent probes: (E) 4.4 kDa Dextran-TRITC, 10 kDA Dextran-AF647 and 70 kDa Dextran-FITC; or (F) 10 kDA Dextran-AF647 and 70 kDa Dextran-FITC and 500 kDa Dextran-TRITC. Early and late swelling phases are highlighted in yellow and green, respectively. (G) Distribution of probe entry times as a function of their hydrodynamic radius. Δ*t* was calculated as the difference between the time of SRB entry (*t* − *t*_0_) and the entry time of the probe of interest. Entry times are shown across the full hydrodynamic radius range (0.5–15.9 nm), showing the rate of pore enlargement. Data are presented as mean *±* SD. Linear fits were performed for SRB, 4.4 and 10 kDa Dextrans (brown line) and for 10, 70 and 500 kDa Dextrans (blue line). For (C,D): *N* =3 experiments, *n*=101 cells; for (E,G): *N* =3, *n*= 49 cells; for (F,G (70kDa and 500 kDA dextran data)): *N* =1, *n*= 31 cells.

The formation of pores in the PM is a defining feature of pyroptosis (10). In this context, GSDMD pores are a prerequisite for triggering cell swelling and the subsequent formation of Ninj1-dependent membrane lesions (9). Structural studies have shown that GSDMD pores can adopt various shapes, including rings, arcs, and slits, which can grow, fuse, and mature (11, 12). GSDMD pores can also dynamically open and close, suggesting that pyroptosis may be reversible (13). However, these findings are based on *in vitro* and *in silico* models, and the precise pore dynamics in live cells remain unknown.

GSDMD pores are non-selective for small solutes, allowing the passage of water and ions (14), as well as fluorescent dyes and proteins such as IL-1*β* (hydrodynamic radius, *R*_H_ = 2.25 nm) (15–18). In contrast, larger molecules such as LDH (*R*_H_ = 4.8 nm) (1) and high-molecular-weight dextrans cannot cross GSDMD pores (1, 3, 17) and instead rely on Ninj1-dependent lesions for their release (3, 8). However, electron microscopy have shown that GSDMD pore size reaches an internal structural radius of ∼ 10 nm within minutes (19), a value largely exceeding the hydrodynamic radius of LDH (*R*_H_ = 4.8 nm). This discrepancy between structural studies and actual transport suggests that passage through GSDMD pores is regulated, possibly via incomplete opening (20) or electrochemical gating (21).

In physiological conditions, the transport across mammalian cell membranes is generally considered to be selective, meaning that ions and water move through the PM via independent routes: ions channels (passive) and ion pumps (active) for ions and aquaporins for water. The integration of such selectivity principle in phenomenological models of water and ions fluxes has been formalized in the Kedem-Katchalsky close-to-equilibrium thermodynamic framework (22, 23). The ensuing relation

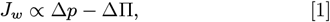

between *J*_*w*_ the water flux and Δ*p* and ΔΠ, the hydrostatic and osmotic pressures difference across the PM is very widely used in cell volume regulation models (24–28).

Relying on relation Eq. (1), the textbook paradigm that allows to understand how a mammalian cell controls its volume is the so called pump and leak model (24, 29). The model allows to maintain the same osmolarity between the cell cytoplasm and the extra-cellular medium by effectively pumping ions out of the cell at a certain rate to counter the passive osmotic effect that tends to homogenize concentrations. Therefore the fact that the cell volume does not keep on swelling in normal situations is related to a non-dimensional parameter comparing the kinetics of active pumping to that of passive leaking.

A correction can be brought by the cell surface and bulk cytoskeleton (30) which also provide an elastic resistance to swelling. However, the typical measured stiffnesses of the cytoskeleton (∼ 0.1 − 1 kPa) are in general negligible compared to that induced by a 10 mM variation of osmotic pressure (∼25 kPa), which is a fluctuation expected to occur in extracellular media that are typically of few hundreds of milimolar. Upon, permeabilization of the PM, one may therefore expect that the passive leaking will completely dominate the active pumping and lead to a very large cell swelling. Since the excess PM surface area stored in membrane invaginations is limited (27, 31), it was widely assumed that the ensuing mechanical stress alone would inevitably rupture the PM - until the discovery of Ninj1 changed this paradigm (3, 9).

Indeed, the historical model suggests that GSDMD pores trigger rapid ion fluxes, osmotic swelling, and mechanical stress culminating in cell lysis (1, 10). This view has since been challenged, first by the identification of Ninj1-dependent PM lesions as a key driver of rupture (3), and then by our recent biophysical study showing that cell swelling unfolds in unexpected two distinct steps not constrained by PM surface availability, indicating that lysis cannot be explained by mechanics alone (9). Together, these findings place PM permeability at the heart of pyroptosis. However, the mechanisms of Ninj1 action, its interplay with GSDMD pores, and the processes linking swelling to rupture remain open questions under active investigation (2, 32).

In this study, we probed the coupling between membrane pore formation and cell swelling dynamics. To this end, we adapted Fluorescence eXclusion microscopy (FXm), a well-established method for the rapid and non-invasive quantification of absolute cell volume in live cells (33– 35) (Fig. 1B & Fig. S1). FXm infers volume from the drop in fluorescence intensity created by a cell due to the exclusion of an impermeable dye. Using a panel of probes of increasing size, we have access to a dual readout of swelling dynamics and pore opening, directly revealed by probe entry. To achieve a comprehensive understanding of these processes, we developed a theoretical framework to model cell volume and PM permeability changes during pyroptosis. Although the PM is rapidly permeabilized to ions, pyroptotic cells progress through successive swelling phases, including a volume plateau, governed by distinct permeability regimes. We demonstrate that GSDMD, Ninj1, and intracellular calcium each leave specific signatures on these phases, establishing the evolution of the pore opening and size as key determinants of swelling dynamics and volume plateau. By developing a simple model to explain these non-trivial dynamics, we emphasize the need to carefully reconsider the commonly assumed selectivity constraints in Eq. (1) when modeling cell volume regulation in lytic cells.

## Results

### Dynamic Correlation of Swelling and PM Permeabilization

Using FXm, we first characterized the pyroptotic cell volume dynamics with high temporal resolution, previously shown to be biphasic (9) in immortalized bone marrow-derived macrophages (WT iBMDMs), a well-established model of pyroptosis. We then validated these findings in mouse embryonic fibroblasts (MEFs). Cells were coincubated with fluorescent probes of increasing hydrodynamic radius (Figs.1 B-D, S2 A-C). The small soluble fluorophore sulfo-rhodamine B (SRB; *R*_H_ = 0.5 nm, blue curve), marks the first PM permeabilization event, while the 10 kDa dextran–AF647 (*R*_H_ = 1.9 nm, black curve) can enter through GSDMD pores, consistent with the reported passage of slightly larger molecules such as IL-1*β* (*R*_H_ = 2.25 nm). In contrast, the much larger 500 kDa dextran (*R*_H_ = 15.9 nm, pink curve Fig.1 D) cannot pass through GSDMD pores as its radius exceeds this size; it allows monitoring of wholecell volume dynamics until its entry upon PM rupture, likely attributed to Ninj1-mediated lesions (3, 9).

The swelling dynamics of pyroptotic cells consistently subdivide into five distinct phases, including the two swelling phases reported previously (9) (Fig.1 D). Pyroptosis begins with a first phase of PM permeabilization, revealed by the rapid cell entry of SRB, which is marked by a strong decrease in SRB-based volume measurements (blue curves, Figs.1 C&D). This permeabilization is correlated with the onset of cell swelling (*p* < 0.001, Fig. S1 A) - here referred to as the “early swelling phase” (phase 2) - detected with 10 and 500 kDa dextrans (Figs. 1 C&D). GSDMD depletion abolishes this early swelling, while inhibition of Na/H exchangers involved in volume regulation had no significant effect (Fig. S2 C&E), indicating that GSDMD pores are essential for the initial phases of PM permeabilization that drives swelling, in line with previous reports (10). At this stage, the PM remains impermeable to molecules larger than 1.9 nm in radius, as 10 kDa dextran does not yet enter. The early swelling phase is typically interrupted by a slowdown, visible as a shoulder in the FXm cell volume curve, which leads to a transient volume plateau in 26% of WT iBMDMs and 40% of MEFs from 1 up to 15 minutes (phase 3, Figs. 1 D, S2 F & S6). We defined a plateau as a period when cell volume remained nearly constant (relative volume variation < 10%) for at least 1 minute. This plateau ends with the entry of 10 kDa dextran, marking the onset of a “late swelling phase” (phase 4) that culminates in PM rupture, detected by 500 kDa dextran entry (phase 5). The two swelling phases are associated with distinct morphological signatures (Fig. S3 D): early swelling corresponds to multiple small blebs, whereas late swelling is characterized by one or two large pyroptotic blebs that expand until PM rupture.

To further resolve the dynamics of PM permeabilization, we applied the same FXm framework using the full panel of fluorescent probes (Fig.1 B) on WT iBMDMs. A sequential entry of dyes is observed according to their size, each entry marked by an increase in intracellular fluorescence and a concomitant drop in the excluded volume (Figs.1 E-F & S3-4). The entry of SRB, marking the first detectable event, served as a temporal reference for evaluating the entry times of the other probes (Fig.1 G). The entry of a given probe indicates that PM pores reached at least the corresponding radius. We computed pore size kinetics by fitting probe entry times to probe size. Fits were performed on two probe groups: SRB, 4.4, and 10 kDa dextrans, since SRB marks the onset of the first swelling phase; and 10, 70, and 500 kDa dextrans, as 10 kDa entry coincides with the onset of the second swelling phase. Two distinct behaviors emerged: (i) the entry of SRB and small dextrans (4.4 and 10 kDa) corresponded to a slow rate of effective pore size (0.29 nm/min), whereas (ii) the entry of larger probes (70 and 500 kDa dextrans) revealed a faster rate of 1.84 nm/min (Fig.1 G). These measurements establish that pyroptotic PM permeabilization proceeds through a gradual enlargement of pores. Importantly, the two distinct rates correspond to early and late swelling, suggesting that these phases arise from different molecular actors. GSDMD which facilitates the initial entry of small dyes, and Ninj1, which acts later to create large lesions enabling passage of larger dyes.

### Ninj1-dependent changes in permeability affect cell swelling dynamics

To dissect the dynamics of pore formation during pyroptosis, we inhibited one of the two key molecular actors, Ninj1, while leaving GSDMD activity intact. Ninj1-dependent PM lesions were blocked either pharmacologically, by treatment with glycine - which prevents Ninj1 oligomerization (38) - or genetically, using *Ninj1*^-/–^ iBMDMs (39) stably transfected with the opto-ASC construct (9). As expected, inhibition of Ninj1 impaired PM rupture, revealed by the altered kinetics of 500 kDa dextran entry (Fig. 2 A-C, pink curve). Glycine treatment delayed 500 kDa dextran entry by ∼ 40 minutes and increased the amount of swelling, with the maximal relative volume at PM rupture reaching around 1.7-fold higher values than in controls (Fig. 2 D–E). In *Ninj1*^-/–^ cells, the phenotype was even more pronounced since no detectable entry of 500 kDa dextrans was observed during the acquisition window, consistent with the presence of GSDMD pores alone (Fig. 2 C). Consequently, it was not possible to quantify either the maximal swelling rate or the time taken for these cells to reach this volume.

**Fig. 2.**
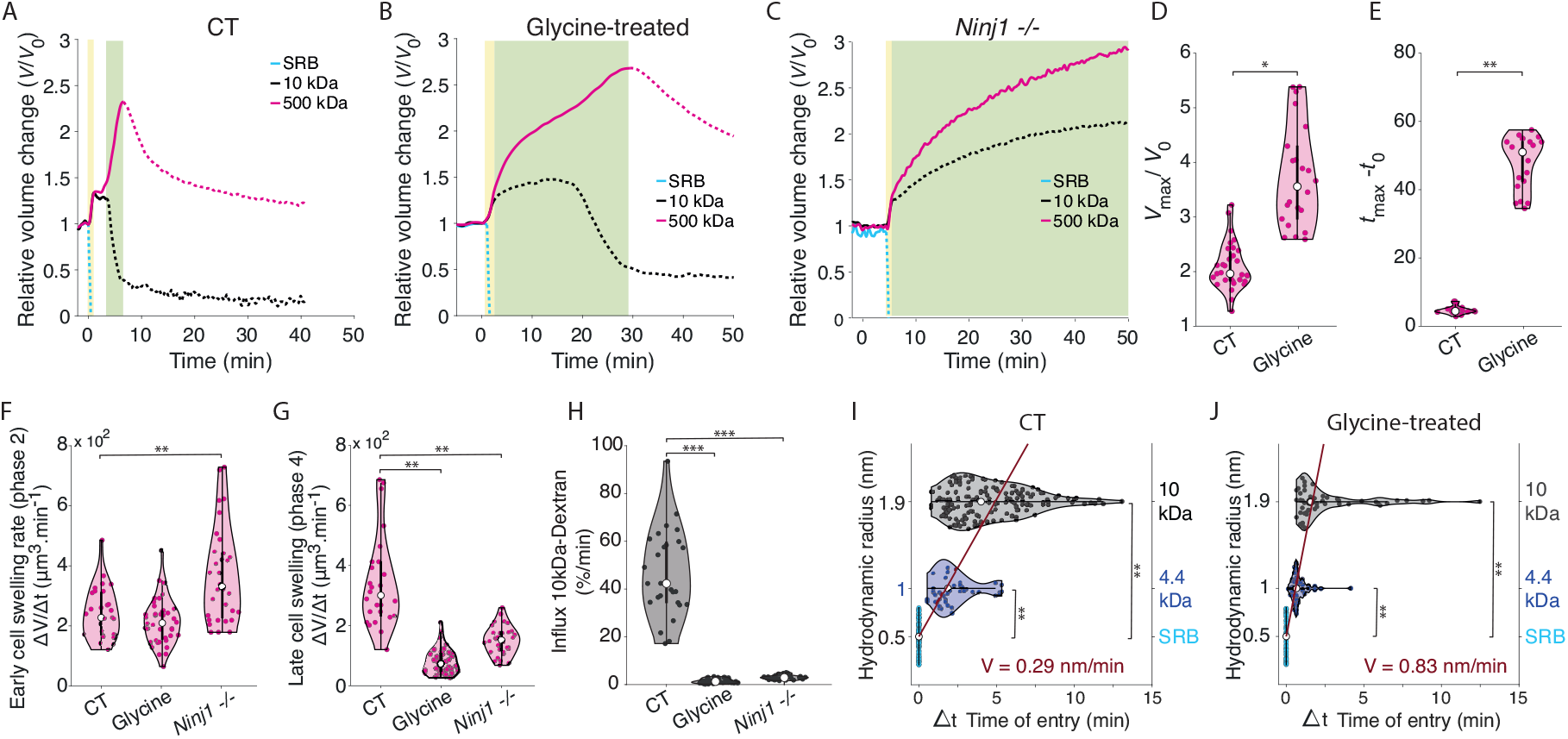
Interplay between Ninj1 and GSDMD pore formation determines pyroptotic swelling dynamics. (A, C, D) Representative normalized dynamics of apparent cell volume during pyroptosis of WT iBMDMs (control (CT), A), WT iBMDMs treated with 20 *µ*M glycine for 2 h (Glycine-treated, B), and *Ninj1*^-/–^ iBMDMs (*Ninj1*^-/–^, C). Dynamics of apparent cell volume were monitored by co-incubation with SRB (blue), 10 kDa Dextran-AF647 (black) and 500 kDa Dextran-FITC (pink). (D) Relative maximal cell volume reached before PM rupture (*V*_max_/*V*_0_, defined by large Dextran entry. (E) Delay between the onset of swelling *t*_0_ and PM rupture *t*_max_. (F) Maximal swelling rate during early swelling phase (pyroptosis phase 2, yellow). (G) Maximal swelling rate during the late swelling phase (pyroptosis phase 4, green). (H) Maximal relative entry rate of 10kDa dextran. (D-G) were computed from 500 kDa dextran excluded volume measurements, and (H) from 10 and 500 kDa Dextran excluded volume measurements. See Materials & Methods for details on parameter computation. (I - J) Entry times of SRB, 4.4 kDa, and 10 kDa Dextrans (hydrodynamic radii 0.5–1.9 nm) in WT iBMDMs in control condition (I) or treated with 20 *µ*M glycine for 2 h (J). Violin plots show the distribution of single-cell data (colored dots), with median (white dot) and SD. Statistical analysis: unpaired t-test (D–H), paired Wilcoxon signed-rank test (J) (\*\**p* < 0.01, \*\**p* < 0.005; *N* =3, *n* =33–103 cells across conditions), and paired t-test (I) (*p* < 0.05; *N* =3, *n* =49 cells).

In addition to impairing PM rupture, inhibition of Ninj1-mediated lesion formation (glycine-treated or *Ninj1*^-/–^) profoundly altered swelling dynamics compared with controls (Fig. 2 A–C). In both conditions, the typical transient volume plateau - separating the early and late swelling phases - was no longer detected. Nevertheless, two distinct swelling phases were identifiable, with distinct solute fluxes (Fig. 2 F&G), consistently separated by the onset of 10 kDa dextran entry (divergence in excluded volumes: 10 kDa *vs* 500 kDa dextran). When Ninj1-mediated lesions are absent in *Ninj1*^-/–^ iBMDMs, the permeability of the early swelling phase is significantly increased, as indicated by the early swelling rate and the SRB influx associated (Fig. 2 F & S5 A). Although glycine treatment had no detectable effect on this stage, it increased the sequential entry rate of SRB and 4.4 and 10 kDa dextrans (0.83 nm/min (Fig. 2 J)) compared with control cells (0.29 nm/min (Fig. 2 I)), indicating a faster enlargement of the GSDMD pore diameter. The absence or inhibition of Ninj1-mediated lesions significantly reduced permeability to 10 kDa dextran (Fig. 2 H) and was associated with a significant decrease in the late swelling rate (Fig. 2 G). Although 10 kDa dextran entered the cell (the excluded volume measured with 10 kDa dextran differing from that measured with 500 kDa dextran (Figs. 2 B-C & S5 F)), its influx was significantly reduced compared with the CT condition confirming that the late swelling phase is mainly Ninj1-dependent. Consistently, entry of 70 kDa dextran was blocked (Figs. S5 D–E), indicating that without Ninj1 the effective GSDMD pore diameter remains close to the 10 kDa cutoff and below the 70 kDa threshold.

### Chelation of intracellular calcium enhances the plateau phase

Intracellular calcium plays multiple roles in pyroptosis, from GSDMD pore formation and Ninj1 activation to PM repair through Endosomal sorting complexes required for transport (ESCRT) proteins (13, 40, 41). Chelation of intracellular calcium did not abolish the biphasic response but strongly delayed the onset of the second swelling phase on iBMDMs (Fig. 3 A). Under intracellular calcium chelation, a higher proportion of cells displayed a volume plateau (Fig. 3 A&B), which was significantly extended compared to the control condition (Fig. 3 D), without affecting its relative value of around 1.5 (Fig. 3 C). Our experimental data shows that the dynamics of cell swelling during pyroptosis is complex, including a slow down of the volume increase after an initial fast swelling. The dynamics of this volume stabilization phase is strongly affected by genetic or pharmacological perturbations (Figs. 2 & 3). In all our experiments, the time of the 10 kDa dextran entry consistently marks the onset of the late swelling phase, in the presence or absence of a plateau. Intracellular calcium chelation or culture in calcium-free medium delayed the 10 kDa dextran entry, whereas Ninj1-lesion inhibition (Glycine-treated WT iBMDMs) accelerated its entry (Figs. 2 J & S7). We also found, in all our tested conditions, a strong inverse correlation between the initial SRB influx and the delay before 10 kDa dextran entry (Fig. S7 B), suggesting that the pore enlargement dynamics is sustained from the early to the late phase.

**Fig. 3.**
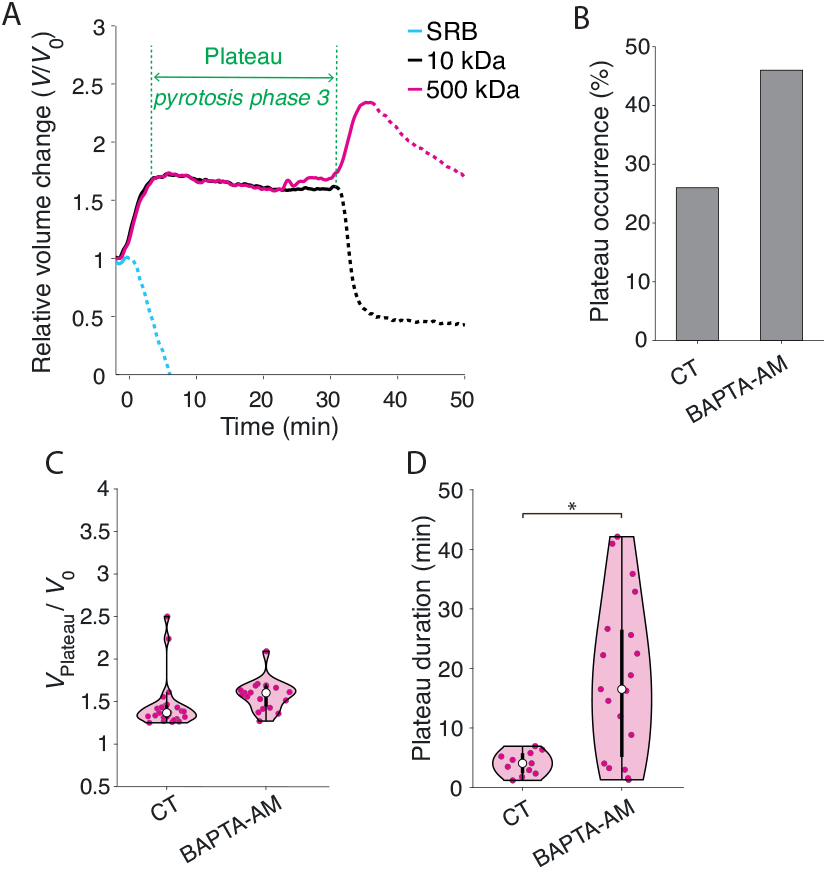
Calcium mediates the plateau phase of pyroptotic swelling. (A) Representative relative curves of apparent cell volume dynamics of WT iBMDMs treated with 20 *µM* BAPTA-AM for 45 min. (B–D) Comparison between untreated (CT) and BAPTA-AM– treated WT iBMDMs of: (B) Frequency of plateau occurrence, (C) relative plateau volume (*V*_p_*/V*_0_), and (D) plateau duration. (C-D) Individual cell data are shown (filled dots) with median (white dot) and SD. Statistical analysis: unpaired t-test (D) *N* = 3, *n* = 87 CT cells and *n* = 31 BAPTA-AM-treated cells. (^*\**^ *p* < 0.05)

Strikingly, the period of volume stabilization occurs despite the plasma membrane remaining permeable to water and ions. In principle, one would expect that a cell with a compromised membrane should swell continuously once ion pumps fail to counter passive osmotic influx. This paradox points to the need for a dedicated theoretical framework to describe cell volume dynamics and membrane fluxes during pyroptosis, raising the question of what physical mechanisms underlie this unexpected stabilization.

### Model for the cell volume control

Before the formation of GSDMD pores, the PM is a highly selective interface (22) and we propose the following simple model for cell volume regulation, only valid at a timescale where synthesis does not increase the cell dry mass in a significant way. Eq. (2) describes the temporal evolution of the volume of water and ion concentration within the cell. See (42) for a detailed thermodynamic close-to-equilibrium derivation of Eq. (2). A scheme of the model is shown on Fig. 4 A:

**Fig. 4.**
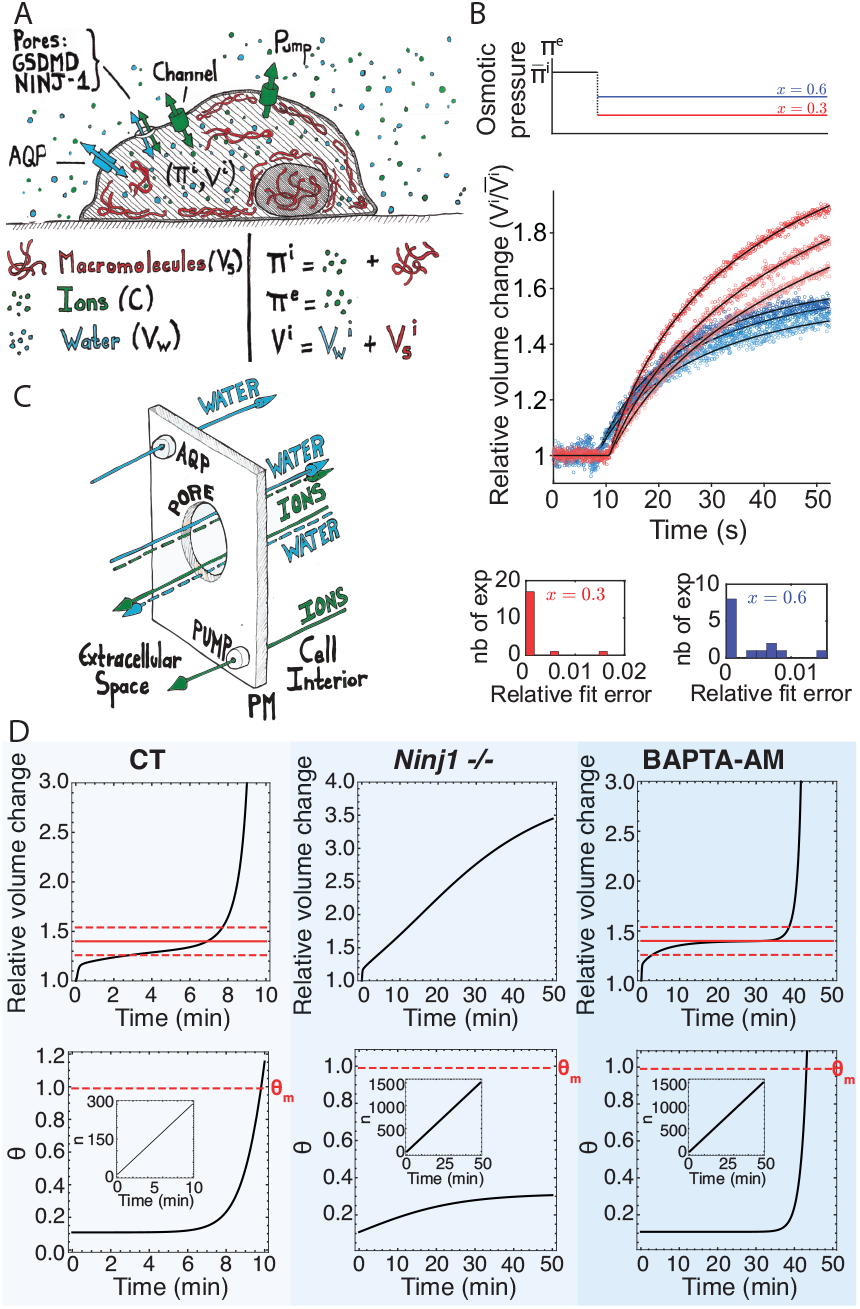
Modeling volume dynamics during pyroptosis. (A) Schematic of the cell volume model. The internal osmotic pressure is caused by the entropy of mixing between macromolecules and ions with water. The external osmotic pressure is only caused by ions. The exchanges at the PM can follow several routes. Ions can travel through channels and pumps while water travels through aquaporins (AQP). GSDMD pores are permeable to both water and ions with a certain selectivity which is assumed to decrease with the pore size. (B) Top, dynamics of osmotic pressure application in WT iBMDMs. Bottom, examples of the experimental volume dynamics and fits according to the model Eq. (5) for WT iBMDMs. The red curves correspond to a *x* = 0.3 dilution factor and the blue *x* = 0.6. The inset plots are bins of the relative squared error between the data and the fit for all the performed experiments in each hypo-osmotic condition (*n* = 14 for *x* = 0.6 and *n* = 19 for *x* = 0.3. See Supplementary Materials & Methods for the fitting parameters. (C) Schematic of modeling of the transport at the plasma membrane at the volume plateau when pores are non-selective for water and ions with coupling between the two fluxes setting the steady state volume. (D) Swelling dynamics obtained with model Eq. (6) with the pore formation dynamics *n* and the evolution of the pore filtration ratio *θ* both given in inserts. The first swelling phase is due to the dynamics of *n* reaching a large value while the second swelling phase is due to the pores getting larger and subsequently becoming less and less selective. The *n* dynamics is taken to be always the same in CT, Ninj1^-/–^ and BAPTA-AM conditions while only an evolution of the pore selectivity parameter *θ* can reproduce the experimentally observed trend of volume swelling dynamics. Parameters are *ϵ* = 10^*− 2*^, *A* = 93, *B* = 3.7, *α* = 1.9 and *τ* = 1s.

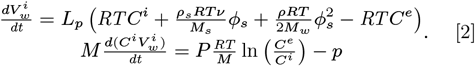

In Eq. (2), 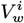 represents the volume of water contained in the cell and *C*^*i*^ is the molar ions concentration in the cell cytoplasm. Water is driven through the PM by the difference of osmotic pressures on both sides with a filtration coefficient *L*_*p*_. The external osmotic pressure *RTC*^*e*^ is, in a dilute approximation, only due to the entropy of mixing of the external ions of concentration *C*^*e*^ where *R* is the gas constant and *T* the temperature. Retaining again only generic entropic terms, the internal osmotic pressure contains three terms. The ions in solution in the solvent lead to the term *RTC*^*i*^, their molar mass is denoted by *M*. The configurational interaction between the water and large macromolecules (43) leads to the two other terms that depend on the cell dry mass volume fraction:

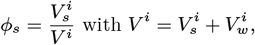

where 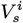 is the fixed volume occupied by the macromolecules and *V* ^*i*^ is the total cell volume (the volume occupied by ions is neglected compared to the volume of water and macromolecules). The *ϕ*_*s*_ term in Eq. (2) is proportional to the molar concentration of macromolecules in the cytoplasm but, in the Flory-Huggins entropy of mixing there is also a 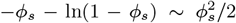 term which is quantitatively important. See (42) for details.

The mass density and the molar mass of water (resp. macromolecules) are denoted *ρ* and *M*_*w*_ (resp. *ρ*_*s*_ and *M*_*s*_). The parameter *? ≃* 0.17 (28) represents the fraction of metabolites among macromolecules since proteins and chromatin contributions are negligible due to their high molar masses (see (42) for details). The mechanical contribution of the PM/cortex complex (Δ*p* term in Eq. (1)) is also neglected in Eq. (2) compared to the osmotic contributions. See (42) for a quantitative justification. The ions move across the PM via selective channels with a permeability *P* and are pumped out at an effective rate *p*.

Introducing the non-dimensional parameters *A, B, α* and *ϵ* and the PM water filtration timescale *τ* defined in Table 1, we can rewrite Eq. (2)

**Table 1.** Rough estimates of the model Eq. (2) parameters. Details and references leading to these effective parameters estimates are given in (42).

| Name | Symbol | Typical value |
| --- | --- | --- |
| membrane permeability | $P$ | $3 * 10^{-19} \text{ g/m}^2 \text{ s}$ |
| ion export rate | $p$ | $5 * 10^{-14} \text{ g/s}$ |
| ion molar mass | $M$ | $30 \text{ g/mol}$ |
| metabolites molar mass | $M_s$ | $200 \text{ g/mol}$ |
| filtration coefficient | $L_p$ | $4 * 10^{-22} \text{ m}^3/\text{s/Pa}$ |
| % of metabolites | $\nu$ | $17\%$ |
| ext. ion concentration | $C^e$ | $300 \text{ mM}$ |
| metabolites mass density | $\rho_s$ | $1300 \text{ g/L}$ |
| water mass density | $\rho_w$ | $1000 \text{ g/L}$ |
| cell radius | $R^i$ | $10 \mu\text{m}$ |
| filtration time | $\tau = V_s^i / (L_p R T C^e)$ | $0.6 \text{ s}$ |
| water mixing | $A = \rho_w / (2 M_w C^e)$ | $93$ |
| metabolites mixing | $B = \rho_s \nu / (M_s C^e)$ | $3.7$ |
| pumping vs leaking | $\alpha = p M / (P R T)$ | $1.96$ |
| ions vs water transport | $\epsilon = P / (L_p (C^e M)^2)$ | $10^{-2}$ |

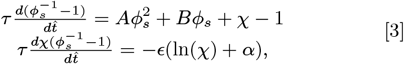

where *χ* = *C*^*i*^*/C*^*e*^ represents the internal ions concentration in the cell compared to the concentration imposed outside. The parameter *A* and *B* represent the osmotic pressure related to the mixing of water and macromolecules compared to the external osmotic pressure of the ions. *α* is the pump-to-leak ratio: the ratio between the rate of active ions pumping compared to the rate of their passive osmotic leaking. Finally *ϵ* controls the permeability of the PM to ions compared to that of water.

This very simple physical model neglects electrostatic effects (28), the differential PM permeabilities of various ions species and the stoichiometry of the pumping mechanisms (24, 44, 45), the various mechanotransduction pathways that affect the PM permeability (25–27) as well as the biochemical regulation of the ion channels and pumps (46). Nevertheless, based on rough experimental data collected in Table 1, the steady state dry mass volume fraction 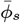 of Eq. (3),

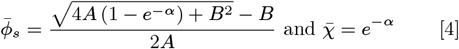

provides the value 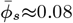. This is similar to the value experimentally found for HeLa cells in (33), 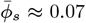. See (42) for details.

To test the model more quantitatively, we performed hypo-osmotic shocks on iBMDMs and MEFs cells by dilution of the external medium and measure their volume with FXm (Fig. 4 B & S8). We only considered a short time scale (< 1 min), therefore probing only the passive volume response leading to water efflux and not active adaptation due to active pumps. This dilution leads to a change of *C*^*e*^ to *xC*^*e*^ where *x* < 1 is the dilution factor. Owing to the fact that ion transport is slow compared to that of water (*ϵ* ≪ 1, see Table 1), we can assume that the number of ions does not change 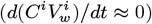 in Eq. (3) leading to the approximation 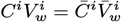 in the tens of seconds following the shock. The bar quantities denote the isotonic situation before the shock.

The cell volume dynamic during the shock is therefore given by the first relation in Eq. (3):

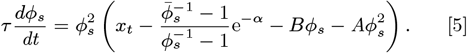

In Eq. (5), *x*_*t*_ switches from the value *x*_*t*_ = 1 when *t* = 0 to *x*_*t*_ = *x* < 1 when *t > t*_*s*_, the time of the shock. The initial condition is 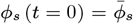. We perform two sets of experiments for the dilution factors *x* = 0.3 and *x* = 0.6.

Such short timescale dynamics is used to estimate the non dimensional parameters *A, B* and *α* and the timescale *τ*. See Fig. 4 B. These values are close to those roughly estimated in Table 1 based on independent literature data, showing that the fitted values are realistic. This confirms that the simple model Eq. (3) is able to quantitatively capture some key features of cell volume regulation at a short timescale.

### Modification of the model in presence of GSDMD pores

As we have already mentioned, a simplistic way to search for a steady state volume in the presence of GSDMD pores would be to consider a vanishingly small pump-to-leak ratio (*α*) in Eq. (4) since GSDMD pores clearly permeabilize the PM to ions (See Fig. 1 D,E,F,G). This however, necessarily leads to a vanishingly small 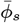 corresponding to an infinite swelling of the cell. An adaptation due to an enhanced pumping rate is very unlikely since the inhibition of Na/H exchanges had no significant effect (Fig. S2 C&E). Modulo a mechanical resistance of the cell membrane which is negligible, this would also happen with more refined models (24–28) which operate with a water flux that follows Eq. (1). However, experimental data (See phase 3 in Fig. 1 D and Fig. 3 A), clearly show the possible existence of a lasting volume plateau after permabilization of the PM by GSDMD pores (See Fig. 3 C&D). We attribute this discrepancy to the fact that the above reasoning considers GSDMD pores in the same way as the selective ion channels. However such pores are also permeable to water, thus also involving cross-coefficients in the transport of water and ions through the PM in this non-selective case. To account for the presence of such coupling, we modify Eq. (2) in the following way (See (42) for the detailed derivation):

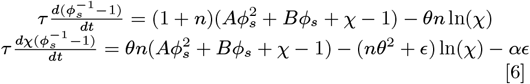

The two additional non-dimensional parameters are *n* representing the permeability to water due to the GSDMD pores in the PM compared to that due to the selective aquaporins and *θ* representing the filtration ratio of the GSDMD pores with respect to the ions. In a caricatural way, we may view *n* as a proxy of the number of GSDMD pores and *θ* as a proxy of the size of these pores, controlling their selectivity. Through both the selective channels and the GSDMD pores, an increase of the ion concentration in the cell will create an influx of water and cell swelling through direct osmosis to dilute the ions. However such increase will also create an efflux of ions to equilibrate the ion concentrations across the PM (Fig. 4 C). This ion flux will create a proportional efflux of water through the pores. The competition between these two phenomena will fix the steady state cell volume. The cross effect also happens symmetrically at the level of ions which flux is no longer only proportional to their difference of concentrations across the PM, but also contains a contribution proportional to the water flux going through the pores and transporting ions. Both direct osmosis and cross-transport are proportional to the number of pores, so the full permeabilization of the PM by the GSDMD pores is not a trivial limit since both terms are still competing.

As a matter of fact, we find that Eq. (6) can still have a finite steady state generalizing Eq. (4) when the permeability to water in the pores is large compare to that of the aquaporins (*n* ≫ 1):

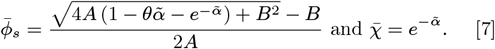

where the effective pump-to-leak ratio reads 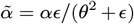. We recover the steady state expressions Eq. (4) when *θ* = 0, which corresponds to a selectivity of the GSDMD pores since only water can get through in this degenerate case. Eq. (7) is only valid in the a range of 0 *≤ θ ≤ θ*_*m*_ where *θ*_*m*_ satisfies:

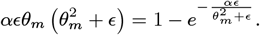

In this range the dry mass volume fraction 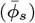 remains finite and is given by Eq. (7) while if *θ > θ*_*m*_, the cell swells to infinity.

Based on these theoretical results, our interpretation of the control case showing two swelling phases of the cell volume separated by a slow down/plateau is the following. The initial formation of GSDMD pores achieves a large permeabilization of the membrane with respect to water and ions. In the model, this can be related to *n* increasing from zero (in the absence of GSDMD pores) to a large value, while *θ* representing the filtration of GSDMD pores with respect to ions, slowly increases to a value *θ*_0_ smaller than the critical value *θ*_*m*_ beyond which no finite swelling is accessible. The consequence is a first swelling phase from the initial homeostatic volume given by Eq. (4) to a shoulder/plateau-like value given by Eq. (7). However,the slow evolution along that dynamics corresponding to the pores getting larger, accelerates close to the end of the plateau. See Fig. 1 G. Such evolution can be captured in the model by a slow and then fast increase of *θ*. The initial dynamics of *n* is important to set the phase 2 swelling velocity. Then *n* takes values that are large enough such that the volume dynamics does not depend on it anymore (corresponding to the limit *n* ≫1). Only the pore selectivity sets the volume dynamics after the phase 2 swelling while the the number of pores is so large that it becomes irrelevant (Fig. 4 D). As *θ* gets towards the critical value *θ*_*m*_, a second swelling phase occurs since no finite volume steady state is any longer accessible. Note that *θ* is an effective parameter in the model representing the level of filtration of the PM by non-selective pores. Such pores are not only represented by GSDMD as Ninj1 also clearly brings a contribution. As a matter of fact, experiments where Ninj1 expression is abolished (Fig. 2 C) can be captured within our model by considering a *θ* parameter dynamic that initially increases faster than in control conditions, suggesting an effect of Ninj1 on the early GSDMD pores opening (Fig. 4 D). This is consistent with our measurements of the pore opening velocity in these conditions. (Fig. 2 I&J). The final brutal increase of *θ* is absent in this case since Ninj1 oligomerization in the PM is no longer present. Within, our model, we can also interpret the chelation of intracellular calcium experiments presented in Fig. 3. Calcium strongly reduces the opening velocity of GSDMD pores and delays the Ninj1 oligomerization in the PM, thus slowing down the increase of *θ* and delaying its final brutal increase due to Ninj1 oligomerization in the PM. In this situation, the saturation of the volume to a plateau value is much more pronounced than in CT conditions (Fig. 4 D).

## Discussion

For long, the mechanisms leading to PM rupture and the release of large molecules during pyroptosis have remained elusive. Recent discoveries of key players, such as GSDMD and Ninj1, have shed light on the underlying molecular mechanisms, significantly enhancing our understanding of the process. Nevertheless, the physical basis of cell swelling and PM rupture remains unclear, as do the precise roles of these players and the chronological sequence of events. In a previous study, we demonstrated that pyroptotic swelling is a two-step process with uncharacterized sequential and molecular events (9). Here, we combined an improved FXm resolution with theoretical modeling and showed that the swelling dynamics can arise from a progressive permeation of the PM by non-selective pores.

Indeed, we showed that cells start to swell as soon as the PM is permeable to small dyes and thus ions. However, even though the PM permeability keeps increasing with the formation of GSDMD pores, this initial swelling slows down and we can even observe a volume plateau that lasts several minutes before another phase of swelling. The plateau is puzzling, as classical models of mammalian cell volume regulation - where active ion efflux by pumps counteracts passive osmotic swelling - cannot account for it (24). In our conditions, as the permeability of the PM is large and the pumps cannot run at an irrealistic pace, such model would predict an infinite cell swelling.

To explain the origin of the volume plateau, we first established the foundation of our model-based on classical pump and leak models - in the absence of pyroptotic pores at the PM (24–28, 44, 45). Our main difference with these models is that we do not consider for simplicity the well-known role of electrostatics in osmosis, but we take into consideration a generic entropy of mixing between biopolymers in solution in the cytosol. Our model can then quantitatively capture the short timescale cell’s passive response to osmotic perturbations with realistic parameters. This demonstrates its ability to quantify key membrane permeability properties, such as the water filtration coefficient and the ion pump-to-leak ratio (Fig. 4 B).

Then, we considered that the pores formed during pyroptosis are non-selective and thus permeable to ions and water since small dyes can pass through. This introduces a coupling between the fluxes of water and ions at the membrane (See Eq. (6) and Fig. 4 C). The competition between water fluxes in the aquaporins and the pores and that of the ions in the pumps/channels and the pores provides the possibility to obtain a volume plateau even in the presence of a large PM permeability. The formation and the non-selectivity of the pores are captured by two parameters in the model, one accounting for the permeability introduced by the pores (*n*) and one for their finite selectivity with respect to ions (*θ*) (See Eq. (6)). Our interpretation of the two-steps swelling is then the following: The first swelling phase is due to the full membrane permeabilization. This phase reaches a volume plateau when the PM permeability due to the pores becomes much larger than that due to the aquaporins. However, this only occurs in a certain range of the pore’s selectivity. The fact that pores effectively become larger (first slowly and then more rapidly, see Fig. 1 G) suggests that the pores become less and less selective over time. As a result, the parameter *θ* is expected to increase beyond the critical threshold after which no plateau volume is anymore possible, as in classical pump and leak models. This is at the origin of the second swelling phase that culminates with the PM rupture.

We further investigated the key molecular players driving changes in permeability (*n*) and selectivity (*θ*). It is well established that pore formation by GSDMD at the plasma membrane is a prerequisite for subsequent rupture mediated by Ninj1 (3, 6, 47), although their dynamics remain under active investigation. We found that the effective size of PM pores gradually increases during pyroptosis, but with two distinct kinetics. A slow, early increase is consistent with the progressive assembly of GSDMD pores, which form within minutes as oligomeric arcs and then slits that already permit ion exchange and water influx (11, 48). Interestingly, by inhibiting Ninj1 activity, we revealed that fully opened GSDMD pores have an effective internal size slightly above 10 kDa dextran but still below 70 kDa, thus providing an upper bound for pore size in living cells.

Subsequently, a more rapid increase rate is attributed to Ninj1-mediated lesion enlargement which are responsible for the large dextran cell entrance. The concept of Ninj1 lesions enlargement is in good agreement with the PM tension-based model for lesion opening proposed recently (32, 39) and the role of Ninj1 in PM fragility (49). In addition, our findings shed light on the requirement of Ninj1-mediated lesion formation for both the establishment of the volume plateau and the onset of the late swelling phase. Indeed, in the absence or inhibition of Ninj1, the plateau disappears and the late phase is strongly altered. Remarkably, this altered volume dynamic coincides with a marked increase in the rate of GSDMD pore opening. This point to an interdependence between GSDMD and Ninj1 which may be mediated by the mechanical properties of the plasma membrane, as no direct interaction has yet been demonstrated. One central actor in pyroptosis is intracellular calcium, whose influx is mediated by GSDMD pores. Calcium has been shown to play a dual role: it can transiently delay plasma membrane lysis by activating ESCRT-dependent membrane repair (41), but it can also promote lysis by driving Ninj1 aggregation into membrane lesions (39, 40). In line with this, we found that calcium chelation prolongs the volume plateau, suggesting that calcium contributes more prominently to Ninj1 activation via lipid scrambling (40) than to ESCRT-mediated membrane repair (41).

Together, these data establish that the dynamics of cell volume during pyroptosis results from the interconnected contributions of GSDMD and Ninj1, each imparting a specific signature on swelling trajectories. The end of the plateau, marking the transition to late swelling, may reflect the formation of large plasma membrane pores mediated by Ninj1, potentially facilitated by intracellular calcium or induced by increased membrane tension due to the complete opening of GSDMD triggering cell swelling (39, 49).

Since most forms of programmed lytic cell death involve significant volume changes and tightly regulated membrane permeability, our approach, combining quantitative volume and permeability tracking with theoretical modeling, offers new insights into both the physical and molecular mechanisms at play. Moreover, comparing different cell death pathways - whether they share volume or permeability dynamics and molecular players or not - can shed light on the underlying physics governing these highly regulated yet dramatic events. For instance, using optogenetic activation, we explored a binary ON/OFF mechanism (9), yet other activators can trigger more graded response for various cell death (50). Capturing the dynamics of gradual activation and healing will definitely provide valuable insights on more subtle events.

## Materials and Methods

### Reagents

Sulfo-Rhodamine B (SRB) and 500 kDa Dextran-FITC were purchased from Sigma-Aldrich. 10 kDa Dextran-Alexa-Fluor 647 (AF647) was purchased from Thermo Fisher. 4.4 kDa Dextran-TRITC, 70 kDa Dextran-FITC, and 500 kDa Dextran-TRITC were purchased from Tdb Labs (Sweden). Polydimethylsiloxane (PDMS), Sylgard 184, was obtained from Dow Corning.

### Cell lines and optogenetic induction of pyroptosis

Murine immortalized bone marrow-derived macrophages (iBMDMs, WT and Ninj1 KO) and mouse embryonic fibroblasts (MEFs) were used. Stable expression of the optogenetic construct opto-ASC (ASC–CRY2–TagRFP) in these lines was achieved as previously described (9). The opto-ASC system, in which light-induced CRY2 oligomerization triggers ASC polymerization, caspase-1 activation, and GSDMD-dependent pyroptosis, faithfully recapitulates classical pyroptotic cell swelling (Fig. 1A). For MEFs, which do not express endogenous caspase-1, stable expression of murine caspase-1 was achieved by lentiviral transduction (9). Cells were cultured in Dulbecco’s Modified Eagle Medium (DMEM, 4.5 g/L glucose; Gibco) supplemented with 10% fetal bovine serum (FBS), 1% penicillin–streptomycin, 1% Glutamax, and 1% sodium pyruvate (all from Invitrogen).

### Live imaging to correlate cell morphology and PM permeability during pyroptosis

Pyroptosis process imaging were performed in a controlled atmosphere (37 °C, 5 % CO2) with a spinning disk confocal microscope Yokogawa CSU-W1 (Objective 40x). Cells were seeded into ibidi µ-slides (VI 0.1), in phenol red-free medium 24 h before the experiment. The medium was then replaced by imaging medium supplemented with 1 g/L of 70 kDa Dextran conjugated to FITC. Optogenetic construct was activated with a 488 nm laser (three times every 10 seconds). Z-stack and timelapse (one image every 15 s) acquisitions were performed over 45 mins to concomitantly visualize speck formation (RFP), bleb appearance, and dextran entry through the PM. Bleb formation was visualized using brightfield microscopy, specks were imaged using a 561 nm laser, and dextran entry coupled to FITC was visualized 488 nm laser. Image analysis was performed with Fiji software.

### Microfluidic chips fabrication for Fluorescence eXclusion microscopy

Microfluidic chips with a fixed chamber height (20.3 µm) – defined by pillar - were made by pouring a mixture of PDMS elastomer and curing agent (1:10, Sylgard 184, Dow Corning) onto a mold - on silicon wafer with SU-8 photoresist made using classical photolithography techniques. The PDMS was then cured for 2 h at 65 °C. After curing, PDMS chips were punched to form inlets and outlets (3 mm of diameter), trimmed, and plasma-bonded to glass coverslips (air plasma, 30 s). Chip/glass coverslip assemblies were mounted on 35 mm Petri dishes using Norland Optical Adhesive NOA 81, allowing full immersion and preventing drying. Finally, chips were UV-sterilized (324 nm, 15 min) and stored in PBS at 4 °C for up to one week before cell seeding.

### Cell Volume measurement using Fluorescence eXclusion microscopy

Cell volume measurements were performed using Fluorescence eXclusion microscopy (FXm) as described previously (33, 35). The day before the experiment, PBS in the chip was replaced by culture medium, and the chips were incubated at 37^*°*^C overnight to allow the saturation of PDMS. 5 *×* 10^4^ cells were seeded in the chamber of microfluidic chips and allowed to adhere overnight. Cells were then incubated with phenol red-free medium supplemented with non-permeable fluorescent dyes and imaging was performed on a Leica DMi8 epifluorescence micro-scope equipped with a 10×/0.3 NA objective, under standard culture conditions (37 °C, 5% CO_2_). Pyroptosis was induced by activating the opto-ASC construct with a 488 nm LED (25% intensity of a CoolLED PE300) for 10 s, delivering a light dose of 3720 mJ/cm^2^. Cell volume was then monitored at an 8-second frame rate using 488, 560, and 635 nm LEDs to visualize FITC-, TRITC-, SRB-, and Alexa Fluor 647–conjugated probes. At cell location, fluorescence is excluded since the dye cannot enter the cell (Fig. S1). Fluorescence was calibrated based on the minimum fluorescence intensity (*I*_min_), measured at the pillar regions, and the maximum intensity (*I*_max_) within the chamber of defined height (*h*_chamber_ = 20.3 µm), thereby defining a linear relationship between fluorescence intensity of a pixel *I*(*x, y*) and the local height of individual cell excluding the fluorescent dye, as established in (33). The volume of each cell was then computed by integrating the fluorescence intensity drop over the cell area:

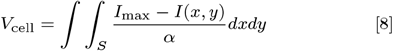

with, *I*_max_ − *I*_min_ = *α ·* (*h*_chamber_), where *α* is the proportionality coefficient. Where, *S* is the cell area and *I*(*x, y*) the local fluorescence intensity.

Volume determination becomes invalid once cells turned permeable to the fluorescent dyes, reflecting PM permeabilization and pore opening dynamics. To accurately monitor the evolution of PM permeability and porosity in relation to cell volume dynamics during pyroptosis, we employed a panel of fluorescent probes with increasing hydrodynamic radii: sulforhodamine B (SRB, *R*_H_ *≈* 0.5 nm), 4.4 kDa Dextran-TRITC (*R*_H_ *≈* 1 nm), 10 kDa Dextran-Alexa Fluor 647 (*R*_H_ *≈* 1.9 nm), 70 kDa Dextran-FITC (*R*_H_ *≈* 6.5 nm), and 500 kDa Dextran-TRITC or conjugated to FITC (*R*_H_ *≈* 15.9 nm) (Fig. 1B). The specific dye sets and experimental conditions used to resolve sequential PM permeabilization steps are described in the supplementary information.

For osmotic-shock experiments, cells were imaged in dedicated low-resistance FXm chambers allowing rapid medium exchange (<1 s) (27). Cell volume was first recorded under isotonic conditions before switching to a hypotonic medium containing Alexa-647–labelled 10 kDa dextran (50 mg/L). Full experimental details are provided in the Supporting Information.

### Image analysis: Cell volume and PM permeability extraction

Image analysis was performed using a homemade MATLAB program adapted from (33, 34). A correction for illumination inhomogeneity was applied to all images. For each time frame and each cell, the linear relation between fluorescence intensity and chamber height was calibrated, ensuring that any global decrease in fluorescence did not affect cell volume measurements (Fig. S1). Cell segmentation was systematically performed on the images of the largest fluorescent dye used, since it is the last to enter the cells. The fluorescence drop was integrated over the segmented cell area and converted into cell volume using Eq. (8). Single-cell volume kinetic curves were obtained for each dye, and characteristic time points were manually extracted, allowing the identification of the five swelling phases occurring during pyroptosis; the criteria used to detect these time points, as well as the quantification of plateau properties and dye influx, are detailed in the supplementary information.

### Statistical analysis

Statistical analysis were performed using MATLAB. Depending on data normality, Two-tailed paired or unpaired Student t-test and non-parametric Mann Whitney test were used, and significance was accepted at the 95 % confidence level (^*\**^ *p* < 0.05, ^*\*\**^, *p* < 0.01, ^*\*\*\**^ *p* < 0.005), with N the number of independent experiments and n the number of analyzed cells.

## ACKNOWLEDGMENTS

We thank Matthieu Mercury from institut Lumière Matière (ILM) for the drawings illustrating the article. Funding: This work was supported by the Projet de Recherche Collaborative (PRC), Agence Nationale de la Recherche (ANR), project SurVol PRC ANR-19-CE13-0030 (to E.B and P.R); Institut Convergence PLAsCAN, ANR-17-CONV-0002 (to E.B., V.P. and S.M.); ETOILE 2022 funding (to E.B. and C.R); Can-céropôle Lyon Auvergne-Rhône-Alpes (to V.P.); Ligue Nationale Contre le Cancer comité de l’Ain (to V.P.); ANR-22-ce15-0032-01 (to V.P).

## Notes

### Competing Interest Statement

The authors have declared no competing interest.

### Summary of Updates

This revised version of the manuscript includes a full update of all figures, of the Results and Discussion sections, of the supplementary data, and we added one coauthor. We incorporated new analyses and new datasets to better distinguish the respective contributions of GSDMD pores and Ninj1 lesions. These data reveal two distinct pore opening dynamics, a slower one driven by GSDMD and a faster one associated with Ninj1. Our analyses also confirm that Ninj1 is essential for the late swelling phase and for plasma membrane rupture, and they show that the absence of Ninj1 alters the kinetics of GSDMD pore opening, indicating an coupling between the two processes. We further demonstrate that both GSDMD pores and Ninj1 lesions expand through gradual pore enlargement, addressing a long-standing question in the field. We also investigated the role of intracellular calcium on volume dynamics. Calcium has little effect on the two swelling phases but prolongs the plateau, suggesting that it contributes to triggering the Ninj1 dependent late swelling phase. These results confirm that calcium is a key modulator of membrane permeability and swelling dynamics during pyroptosis. In addition, we strengthened the description of our physical model. The revised model now provides a simple and robust framework that accounts for the nonlinear swelling dynamics observed experimentally. It captures how the sequential actions of GSDMD and Ninj1 define permeability regimes and volume responses, and it reproduces major features of the data, including the plateau phase, its modulation by calcium, and its loss in the absence of Ninj1 lesions. This integrated modeling and experimental approach offers a unified biophysical interpretation of pyroptotic volume changes.

