## Supplemental Information for "Pore-size dynamics control complex volume swelling in pyroptosis"

This supplementary information presents the experimental methods complementing the main text, the details of the derivation and analysis of our theoretical model as well as figures reporting some experimental data complementing the main text.

### EXTENDED METHODS

#### Cell Volume measurement using Fluorescence eXclusion microscopy

Fluorescent probe combinations and experimental conditions used to monitor sequential PM permeabilization steps are detailed below. Cells were first incubated with SRB (0.1 g/L), 10 kDa Dextran-Alexa Fluor 647 (50 mg/L), and 500 kDa Dextran-FITC (0.5 g/L) to simultaneously monitor early PM permeabilization (SRB entry), selective passage through GSDMD pores ( 10.75 nm radius; 10 kDa entry), and terminal PM rupture mediated by Ninj1 (500 kDa entry). Global cell volume variations were quantified from 500 kDa Dextran–based excluded volume measurements up to permeabilization. These experiments were performed under control conditions on WT iBMDMs and MEFs, as well as on iBMDMs pre-treated for 45 min with 20 µM BAPTA-AM and maintained during acquisition with 20 µM BAPTA-AM and 1 mM probenecid, on WT iBMDMs treated with 20 µM glycine for 2 h, and on Ninj1-deficient iBMDMs (*Ninj1*$^{-/-}$).

To further resolve PM pore dynamics, WT iBMDMs and MEFs cells were incubated either with SRB (0.1 g/L), 4.4 kDa Dextran-TRITC (1 g/L), and 10 kDa Dextran-Alexa Fluor 647 (50 mg/L) to follow progressive GSDMD pore opening, or with 10 kDa Dextran-Alexa Fluor 647 (50 mg/L), 70 kDa Dextran-FITC (2 g/L), and 500 kDa Dextran-TRITC (1 g/L) to capture larger Ninj1-dependent lesions. The 10 kDa Dextran was included in all conditions as a reference marker to ensure consistency across experiments.

For osmotic shocks, the surface of the microfluidic chambers was coated right after plasma bonding with 0.01% poly-L-lysine solution for 30 minutes (Sigma-Aldrich). Cells were injected and allowed to adhere for 15 minutes before injection of the fluorescent dye-containing isotonic medium (Alexa-647 10 kDa Dextran 50 mg/L). Cell volume monitoring using FXm was initiated under isotonic conditions, after which the medium was exchanged for a known hypotonic solution supplemented with Alexa-647 10 kDa Dextran (50 mg/L). The whole process was performed during continuous FXm acquisition with a 100 ms frame rate using 635 nm LED to visualize Alexa Fluor 647 fluorescence. The hypo-osmotic solution was prepared by adding water to the culture medium.

#### Image analysis: Cell volume and PM permeability extraction

This section details the criteria used to detect characteristic time points during pyroptosis, as well as the procedures used to quantify volume plateau properties and dye influx kinetics from FXm measurements. The FXm signal with SRB incubation was used to detect the first permeabilization phase (onset of cell swelling), corresponding to the earliest detectable PM permeability. This time point was named $t_0$, with corresponding cell volume $V_0$; both values were used to normalize cell volume curves.

All subsequent cell volume changes during pyroptosis were tracked based on the FXm signal obtained with 500 kDa-Dextran–FITC, which has the largest hydrodynamic diameter. The early swelling phase (phase 2) was defined starting at $t_0$, marked by the initial increase in cell volume until stabilization. The end of phase 2 was determined at time $t_{\mathrm{p,start}}$, with corresponding cell volume $V_{\mathrm{p,start}}$, identified in all cases by an inflection of the volume curve and, in some cases, by a cell volume plateau until $t_{\mathrm{p,end}}$, corresponding to cell volume $V_{\mathrm{p,end}}$. The cell volume plateau defined phase 3. The plateau was identified when cell volume stabilized during swelling according to the condition

$$\frac{V_{\mathrm{p,end}} - V_{\mathrm{p,start}}}{V_{\mathrm{i}}} < 10\% \quad \text{and} \quad t_{\mathrm{p,end}} - t_{\mathrm{p,start}} > 1 \text{ min}.$$

The relative plateau volume change $\Delta V_{\mathrm{p}}/V_0$ was then calculated from $V_{\mathrm{p,start}}/V_0$.

The end of phase 3 ($t_{\mathrm{p,end}}$, $V_{\mathrm{p,end}}$) defined the start of the late swelling phase (phase 4), which ended with 500 kDa-Dextran–FITC entry. This event was characterized by a fluorescence increase within the segmented cell area.

From this time point, cell volume was no longer assessed. The 500 kDa-Dextran–FITC entry determined PM rupture and the maximum cell volume ($V_{\max}$) at $t_{\max}$.

The rates of early (phase 2) and late (phase 4) swelling were determined by linear fitting of the excluded cell volume measured with 500 kDa dextran. Fits were initiated at the onset of each swelling phase (at $V_0$ and at the time of 10 kDa entry, $t_{10,\text{in}}$, respectively), over the linear region comprising at least 7 consecutive points, and the slope was taken as the swelling rate.

For each Dextran, the delay $\Delta t$ between SRB entry ($t_0$) and its entry into the cell (detected as a fluorescence increase within the segmented cell area) was calculated to determine their sequential entry into the cytoplasm.

Influx of 10 kDa dextran was quantified by calculating the relative difference between excluded volumes measured with 10 kDa and 500 kDa dextran, relative to the excluded volume obtained with 500 kDa dextran at the time of 10 kDa entry ($t_{10,\text{ in}}$, corresponding to the beginning of phase 4, See Fig. S7F): $100 \cdot (V_{10}(t) - V_{500}(t))/V_{500}(t_{10,\text{ in}})$ where $V_{10}$ and $V_{500}$, are excluded volumes measured with 10 kDa and 500 kDa dextran, respectively. Then linear fittings ($f = -\alpha \cdot t$) were performed locally on the relative difference curve with a moving window of 7 time points. The mean of the absolute values of the top 20% slope values ($\alpha$) was computed.

To obtain the relative SBR influx, a similar approach was used. The relative excluded volume of SRB was computed ($100 \cdot V_{SRB}(t)/V_{SRB}(t_0)$) and the absolute mean of the top 10% slope values was computed.

### Fitting parameters for osmotic shocks

For WT iBMDMs cells, we find the fitting parameters $A = 92.50 \pm 1.23$, $B = 4.40 \pm 0.65$, $\alpha = 2.07 \pm 0.36$ and $\tau = 1.09 \pm 0.28$ s for $x = 0.6$ and $A = 93.35 \pm 1.35$, $B = 3.79 \pm 0.63$, $\alpha = 2.26 \pm 0.35$ and $\tau = 0.99 \pm 0.26$s for $x = 0.3$.

For MEF cells, we find the fitting parameters $A = 75.73 \pm 23.55$, $B = 7.50 \pm 4.27$, $\alpha = 5.18 \pm 1.82$ and $\tau = 0.43 \pm 0.32$ s for $x = 0.6$ and $A = 88.22 \pm 13.34$, $B = 3.97 \pm 3.83$, $\alpha = 6.76 \pm 2.02$ and $\tau = 0.56 \pm 0.50$s for $x = 0.3$.

### A MODEL FOR THE CELL VOLUME PLATEAU IN THE PRESENCE OF GSDM PORES

We consider a system consisting of a cell of volume $V^i$ in an external medium of fluid of volume $V^e$. The cell is composed of a scaffold of large macromolecules (typically proteins, lipids, metabolites, nucleic acids, etc) and small diffusible species (typically ions). At our timescale of interest, only water and ions can move through the cell plasma membrane while larger macromolecules are trapped inside the cytoplasm.

We derive from linear thermodynamics a simple cell volume regulation model that effectively accounts for ions leaking in and out the cell membrane through osmosis and being actively pumped out of the cell [16]. The presence of more ionic species, their complex interactions and their differential membrane permeabilities [13] and pumping rates are not taken into account in our effective model. We also neglect for the sake of simplicity and transparency the electrostatic effects between the ions and the macromolecules, which can bring an important contribution to the osmotic pressure [14, 17, 19, 23]. However, we consider the presence of large macromolecules and their osmotic interaction with the solvent, which is seldom the case in the context of cell volume regulation. The aim of the model is to show how the volume of the cell can stabilize to a plateau due to the formation of non-selective GSDMD pores within the cell membrane.

### I. CONSERVATION LAWS

The balance of the mass of water across the cell membrane reads

$$\rho_w \frac{dV_w^i}{dt} = J_w = -\rho_w \frac{dV_w^e}{dt}, \tag{1}$$

where $\rho_w$ is the mass density of water (the solvent of the cytosol), $V_w^i$ is the volume of water in the cell and $V_w^e$ is the volume of water out of the cell. The flux of water at the cell membrane is denoted $J_w$. Similarly, the mass balance of large macromolecules trapped inside the cell is

$$\rho_s \frac{dV_s^i}{dt} = 0,$$

where $\rho_s$ is the mass density of macromolecules and $V_s^i$ is the dry cell volume. As synthesis is not considered at our timescale of interest, $V_s^i$ is a constant quantity. We finally consider the mass fraction $\phi^i$ of ions in solution in the cell.

The mass fraction of ions outside is denoted $\phi^e$. The mass fraction is the mass of the ions divided by the mass of water in a representative volume element. It is assumed to be a small quantity. The mass balance laws for the ions are

$$\rho_w \frac{d(\phi^i V_w^i)}{dt} = S = -\rho_w \frac{d(\phi^e V_w^e)}{dt}. \tag{2}$$

The ion flux at the plasma membrane is denoted $S$. The total cell volume is

$$V^i = V_w^i + V_s^i, \tag{3}$$

since we neglect the volume occupied by ions in the cell as the dry mass of ions occupies about 3% of the total dry mass of the cell [27]. We thus need to find $V_w^i$ (the other contribution being fixed) by specifying the membrane fluxes $J_w$ and $S$. We do so in the next section following a close-to-equilibrium thermodynamic procedure.

### II. THERMODYNAMICS

The mechanical power applied on the system is

$$\frac{dW}{dt} = -p^i \frac{dV_w^i}{dt} - p^e \frac{dV_w^e}{dt},$$

where we only consider the hydrostatic part of the mechanical stress and $p^i$ and $p^e$ are the internal and external pressures in the incompressible fluid phases.

The system is subjected to some energy input from the external medium. In line with the active gel theory [20, 26], we model this metabolic part as an external power that depends on the the extent of reaction $\zeta$ of the ATP hydrolysis with a fixed affinity $A$:

$$\frac{dP_{\text{ext}}}{dt} = A \frac{d\zeta}{dt}.$$

Next, we suppose that the free energy of the system can be split in the following contributions [4]:

$$F = \rho_s V_s^i f_s(\phi_s) + \rho_w V_w^i f_i\left(\phi^i, \phi_s\right) + \rho_w V_w^e f_e\left(\phi^e\right).$$

The dependence of the first two terms on the volume fraction of the macromolecules in the cell

$$\phi_s = V_s^i / V^i$$

accounts for the mixing of the macromolecules and the solvent [9]. The $\phi^{i,e}$ dependencies represent the osmotic interaction between the ions and the solvent.

Assuming that the system remains at a constant temperature, the dissipation

$$D = \frac{dW}{dt} + \frac{dP_{\text{ext}}}{dt} - \frac{dF}{dt},$$

has to remain non-negative according to the second principle. Using the conservation laws (1)-(2), we obtain

$$\frac{dF}{dt} = J_w(f_i - f_e) + \mu^i\left(S - J_w\phi^i\right) - \mu^e\left(S - J_w\phi^e\right) + J_w\left[\mu_s^i(\phi_s^2 - \phi_s) - \mu_s^s\frac{\rho_s}{\rho_w}\phi_s^2\right],$$

where the chemical potentials are defined in the classical way: $\mu^{i,e} = \partial f_{i,e}/\partial\phi^{i,e}$, $\mu_s^i = \partial f_i/\partial\phi_s$ and $\mu_s^s = \partial f_s/\partial\phi_s$. We therefore obtain the following expression for the dissipation

$$D = J_w\left(\Delta p - \Delta\Pi\right)/\rho_w + S\Delta\mu + Ad\zeta/dt \geq 0,$$

where $\Delta h = h^e - h^i$ denotes the difference of the considered quantity $h$ between the extracellular medium and the intracellular medium and the osmotic pressures are defined as:

$$\Pi^i = \rho_w(\phi^i\mu^i - f_i + \mu_s^i\left(\phi_s - \phi_s^2\right)) + \rho_s\mu_s^s\phi_s^2 \text{ and } \Pi^e = \rho_w(\phi^e\mu^e - f_e).$$

Close to thermodynamic equilibrium, following the Onsager principle, generalized fluxes can be related to generalized forces through a matrix $\Lambda$ of kinetic coefficients [7]:

$$\begin{pmatrix} J_w/\rho_w \\ S \\ d\zeta/dt \end{pmatrix} = \begin{bmatrix} \Lambda_{11} & \Lambda_{12} & \Lambda_{13} \\ \Lambda_{21} & \Lambda_{22} & \Lambda_{23} \\ \Lambda_{31} & \Lambda_{32} & \Lambda_{33} \end{bmatrix} \begin{pmatrix} \Delta p - \Delta\Pi \\ \Delta\mu \\ A \end{pmatrix}. \tag{4}$$

Next, we specify the form of the free energies used to compute the generalized fluxes.

### III. CONSTITUTIVE ASSUMPTIONS AND SIMPLIFICATIONS

We consider the generic case where the free energies only contain the classical entropy of mixing terms:

$$f_i = \frac{RT}{M}(\phi^i \log \phi^i - \phi^i) + \frac{RT}{M_w} \log (1 - \phi_s) \, , f_e = \frac{RT}{M}(\phi^e \log \phi^e - \phi^e), f_s = \frac{RT}{M_s} \log(\phi_s),$$

where $R$ is the gas constant, $M$ is the molar mass of the ion, $M_w$ is the molar mass of water and $M_s$ is the molar mass of macromolecules. Using the above expressions of the free energies, the chemical potential across the membrane takes the usual form:

$$\Delta\mu = \frac{RT}{M} \log \left( \frac{C^e}{C^i} \right) \tag{5}$$

and that of osmotic pressure is:

$$\Delta\Pi = RT \left( C^e - C^i \right) + \frac{\rho_w RT}{M_w}(\phi_s + \log(1 - \phi_s)) - \frac{\rho_s RT}{M_s}\phi_s \approx RT \left( C^e - C^i \right) - \frac{\rho_w RT}{2M_w}\phi_s^2 - \frac{\rho_s RT}{M_s}\phi_s,$$

where the molar concentrations are defined as $C^{i,e} = \rho_w \phi^{i,e}/M$. The last approximation relies on measurements performed on HeLa cells [33] for which the mean volume during the cell cycle is $V^i \approx 2700 \mu\text{m}^3$ and the mean dry mass is $\rho_s V_s^i \approx 255$pg. Taking for $\rho_s \approx$ 1300 g/L (based on the rough estimates of the mass densities of chromatin 1700 g/L, lipids 950 g/L, proteins 1300 g/L and sugars 1500 g/L), the approximate mass density of macromolecules, we obtain $\phi_s \approx 0.07$. Thus $\log(1 - \phi_s) + \phi_s \approx -\phi_s^2/2$ and we denote $\Pi^s$ the contribution to the osmotic pressure due to the interaction between the solvent and the macromolecules:

$$\Pi^s = \frac{\rho_s RT}{M_s}\phi_s + \frac{\rho_w RT}{2M_w}\phi_s^2. \tag{6}$$

To estimate the order of magnitude of the two terms entering in (6), we follow [27] and separate the cell dry mass trapped within the cell membrane into three categories: the proteins, the metabolites and the chromatin. The proportions of the dry mass of these three classes of macromolecules are estimated in [27] and references therein, 60% for the proteins, 17% for the metabolites and 7% for the chromatin. Due to the relative homogeneity between the mass density of all of these species and that of water, we infer that these proportions also approximate the contributions to the volume fraction $\phi_s$. Each of these categories then contributes to the first term of $\Pi^s$ in (6) in a different way.

- The contribution of the proteins can be estimated to

  $$\frac{\rho_s RT}{M_s}\phi_s \approx \frac{1300 \times 8.31 \times 300}{60000}(0.07 \times 0.6) \approx 2 \text{ kPa}.$$

- That of metabolites is much more important with,

  $$\frac{\rho_s RT}{M_s}\phi_s \approx \frac{1300 \times 8.31 \times 300}{200}(0.07 \times 0.17) \approx 190 \text{ kPa}.$$

- And finally the contribution of chromatin is completely negligible since:

  $$\frac{\rho_s RT}{M_s}\phi_s \approx \frac{1300 \times 8.31 \times 300}{12 \times 10^6 \times 650/32}(0.07 \times 0.07) \approx 6 \times 10^{-5} \text{ kPa}.$$

  The molar mass of chromatin is determined based on that of budding yeast [27]. The budding yeast genome contains 12M base pairs with a molar mass of roughly 650 g/mol separated in 16 pairs of chromosomes.

Only the term related to the metabolites is within the same range as the term due to the solvent:

$$\frac{\rho_w RT}{2M_w}\phi_s^2 \approx \frac{1000 \times 8.31 \times 300}{2 \times 18} 0.07^2 \approx 340 \text{ kPa}.$$

In the following, we therefore approximate the osmotic pressure due to the configurational interaction between the the solvent and the macromolecules by the contribution:

$$\Pi^s = \pi_s \phi_s + \pi_w \phi_s^2.$$

| name | symbol | typical value |
|---|---|---|
| membrane permeability | $P$ | $3 \times 10^{-19}$g/m$^2 s$ |
| ion export rate | $p$ | $5 \times 10^{-14}$ g/s |
| ion molar mass | $M$ | 30 g/mol |
| metabolites molar mass | $M_s$ | 200 g/mol |
| membrane filtration coefficient | $L_p$ | $4 \times 10^{-22}$m$^3$/s/Pa |
| proportion of metabolites | $\nu$ | 17% |
| external ion concentration | $C^e$ | 300 mM |
| mass density of metabolites | $\rho_s$ | 1300g/L |
| mass density of water | $\rho_w$ | 1000g/L |
| cell radius | $R^i$ | 10$\mu$m |
| filtration time | $\tau = V_s^i/(L_p RTC^e)$ | 0.6 s |
| water mixing | $A = \rho_w/(2M_w C^e)$ | 93 |
| metabolites mixing | $B = \rho_s \nu/(M_s C^e)$ | 3.7 |
| pumping versus leaking ratio | $\alpha = pM/(PRT)$ | 1.96 |
| ions vs water transport ratio | $\epsilon = P/(L_p(C^e M)^2)$ | $10^{-2}$ |

TABLE I. Rough estimates of the model (9) parameters. A typical value of the membrane conductivity is given in [19]: $g = 3 \times 10^{-9}$ C/V/s, assuming the cell surface area $S_i = 4\pi(R^i)^2 \approx 10^{-9}$m$^2$. The membrane ion permeability is related to the conductivity through $P = gM^2/\mathcal{F}^2$ where $\mathcal{F} = 9.6485 \times 10^4$ C/mol is the Faraday constant. We therefore obtain an effective value for $P$ of the order of $P \approx 3 \times 10^{-19}$g/m$^2 s$ with $M \approx 30$ g/mol the typical molar mass of ions present in the cell. A characteristic value of the membrane filtration coefficient is given in [8], where the value per unit surface $l_p = 4 \times 10^{-13}$ m/s/Pa can be multiplied by the estimate of the total cell surface $S_i$, leading to $L_p = 4 \times 10^{-22}$m$^3$/s/Pa in agreement with the value $4.4 \times 10^{-22}$m$^3$/s/Pa given in [2]. Thus we obtain that $\epsilon \approx 10^{-2}$. We estimate $V_s^i \simeq 2 \times 10^{-16}$m$^3$ based on [33] (see Sec. III) and $RTC^e$≈750kPa. This leads to the rough estimate $\tau \simeq 0.6$s. Note that when we consider the exchange rates at the membrane, we take a fixed approximate effective radius $R^i$ and do not consider the effect of a change of $R^i$ on the available exchange surface area. This is because as the volume increases and the membrane is getting stretched, the exchange surface does not necessarily increases since the membrane contains a large amount of folds and structures that can be released under tension. It is estimated that the excess of membrane surface area is about three times the apparent surface based on a spherical assumption (see [31] and references therein) but it is however unclear how much of this stored surface participates in ions exchanges. Based on the order of magnitudes discussed in Sec. III, we find that $A$≈93 and $B$≈3.7. $\alpha$ is estimated in [27] to be $\alpha \approx 1.96$. A similar estimate of $\alpha \approx 1.6$ is given in [19]. With the above value of membrane conductivity and the estimate of the total cell surface $S_i$, this corresponds to an effective ion pumping rate of $10^9$ ions per second for the whole cell. Such pumping rate value can be compared with the estimate of the number of cycles ≈ 100 that a Na/K ATPase pump performs in one second [11] times the density of Na/K ATPase pumps ≈ 1000$\mu$m$^{-2}$ per unit surface membrane [6, 25] times the total cell surface, leading to the order of magnitude of of $10^8$ ions per second for the whole cell, in the same ballpark.

In the above formula $\pi_s = \rho_s RT\nu/M_s$ with $\nu \approx 0.17$ the proportion of metabolites in the cell and $M_s$ the approximate molar mass of metabolites ($M_s$≈200 g/mol) and $\pi_w = \rho_w RT/(2M_w)$. This osmotic pressure term due to the mixing between the solvent and the macromolecules has sometimes been a bit overlooked with the argument that the concentration of macromolecules in number per unit volume is very small compared to that of ions [14]. Based on the value of an external ions concentration of $C^e$≈300 mM [2, 22], the ion osmotic external pressure can be estimated as

$$\Pi^e = RTC^e \approx 750\text{kPa}.$$

This value is much larger than the osmotic pressure induced by proteins or chromatin but is is within the same range as the osmotic pressure induced by metabolites. The second term in (6) due to the solvent is therefore also important.

In the models [14, 17, 19, 23, 27], the $\Pi_s$ term is neglected but the role of the macromolecules is still present as they lead to an indirect ions-mediated osmotic effect. This is because the macromolecules carry an important number of negative charges that need to be neutralized by their counterions. We deliberately neglect these effects in our simple model to show that another physical ingredient can also lead to a quantitative model of the cell volume regulation.

Finally, assuming a spherical symmetry, the force balance at the cell membrane/cortex can be written

$$\Delta p = -\frac{2\gamma}{R^i} \ ,$$

where $R^i = \sqrt[3]{3V^i/4\pi}$ is the cell radius and $\gamma$ is the tension in the membrane/cortex of the cell [5]. A rough estimate of such tension is given in [30]: $\gamma \approx 4 \times 10^{-4}$ N/m. With the typical value $R^i \approx 10\mu$m, we obtain $\Delta p$≈100 Pa. This jump of hydrostatic pressure is therefore negligible compared to the typical values of the ion osmotic external pressure $\Pi^e \approx 750$ kPa.

In the context of cell volume regulation, the generalized thermodynamic force responsible the water flow in and out the cell can thus be approximated by its osmotic contribution:

$$\Delta p - \Delta\Pi \approx -RT\left(C^e - C^i\right) + \Pi^s$$

With these assumptions, the volume dynamic of the cell can be determined as soon as the permeability matrix coefficients $\Lambda$ in (4) are known.

### IV. STEADY STATE VOLUME IN NORMAL CONDITIONS

In normal conditions, the cell membrane is a highly selective interface [18] such that $\Lambda$ is of the following form

$$\begin{pmatrix} J_w/\rho_w \\ S \\ d\zeta/dt \end{pmatrix} = \begin{bmatrix} L_p & 0 & 0 \\ 0 & P & \Lambda_{23} \\ 0 & -\Lambda_{23} & \lambda \end{bmatrix} \begin{pmatrix} \Delta p - \Delta\Pi \\ \Delta\mu \\ A \end{pmatrix}, \tag{7}$$

where we have considered that aquaporins and ion channels are selective (i.e. ions and water travel through the membrane following separate routes) and that the ion flux at the membrane also effectively includes an active term representing the ATP-driven ion pumps. The filtration coefficient of the membrane is denoted $L_p$ and the membrane permeability to ions is $P$. The cross-coefficient $\Lambda_{23}$ is related to the pumping rate and fixes the export rate $p = -\Lambda_{23}A \geq 0$. We have also assumed that $\Lambda_{32} = -\Lambda_{23}$. The situation is similar in the active gel theory [20] where the active stress mediated by the molecular motors using chemical energy appears as a reactive flux. This is based on the fact that $\Delta p - \Delta\Pi$ and $\Delta\mu$ are odd under time reversal while $A$ is even.

Using the expressions (5)-(6), we therefore obtain the coupled system modeling the dynamics of the cell volume $V^i = V^i_w + V^i_s$ :

$$\begin{cases} \frac{dV^i_w}{dt} = L_p\left(-RT\left(C^e - C^i\right) + \pi_s\phi_s + \pi_w\phi_s^2\right) \\ M\frac{d(C^iV^i_w)}{dt} = P\frac{RT}{M}\log\left(\frac{C^e}{C^i}\right) - p, \end{cases} \tag{8}$$

where $\phi_s = V^i_s/(V^i_w + V^i_s)$ and we recall that $V^i_s$ is a fixed quantity since no synthesis is considered at our timescale of interest. Introducing the non-dimensional parameters:

$$A = \frac{\pi_w}{RTC^e},\ B = \frac{\pi_s}{RTC^e},\ \alpha = \frac{pM}{PRT} \text{ and } \epsilon = \frac{P}{L_p(C^eM)^2},$$

we obtain,

$$\begin{cases} \tau\frac{d(1/\phi_s - 1)}{d\hat{t}} = A\phi_s^2 + B\phi_s + \chi - 1 \\ \frac{\tau}{\epsilon}\frac{d\chi(1/\phi_s - 1)}{d\hat{t}} = -(\log(\chi) + \alpha) \end{cases} \tag{9}$$

where $\chi = C^i/C^e$ represents the internal ions concentration in the cell compared to the concentration imposed outside and the reference timescale is

$$\tau = \frac{V^i_s}{L_pRTC^e}.$$

The parameters $A$ and $B$ represent the osmotic pressure of the mixing of water and macromolecules compared to the external pressure of the ions. The parameter $\alpha$ is a ratio between the rate of ions pumping compared to the rate of their passive osmotic leaking. Finally, the non dimensional parameter $\epsilon$ controls the permeability of the membrane to ions compared to that of water. These non-dimensional parameters are estimated in Table I.

The steady state dry mass volume fraction $\bar{\phi}_s$ corresponding to $J_w = S = 0$ in (9) can then be determined by solving,

$$A\bar{\phi}_s^2 + B\bar{\phi}_s + e^{-\alpha} - 1 = 0. \tag{10}$$

The positive solution of (10) is

$$\bar{\phi}_s = \frac{\sqrt{B^2 + 4A(1 - \mathrm{e}^{-\alpha})} - B}{2A} \Rightarrow \bar{V}^i = \frac{V_s^i}{\bar{\phi}_s},\ \bar{V}_w^i = V_s^i(\frac{1}{\bar{\phi}_s} - 1) \text{ and } \bar{C}^i = C^e \mathrm{e}^{-\alpha} \tag{11}$$

With the parameters estimates of Table I, (11) provides the value $\bar{\phi}_s \approx 0.078$, very close to the value experimentally found in [33]. Even if the present model is extremely simple as it neglects electrostatic effects [27], the differential membrane permeabilities of various ions species and the stoichiometry of the pumping mechanisms [19, 23] as well as the biochemical regulation of the ion channels and pumps [22], it nevertheless provides a reasonable estimate for $\bar{\phi}_s$.

If the active parameter vanishes ($\alpha = 0$), $\bar{\phi}_s = 0$ which means that the cell volume swells to infinity. The work of the pumps is necessary to maintain a steady state cell volume. This effect is recognized to be crucial in the regulation of mammalian cells' volume [13]. The limit of a small pumping rate $\alpha \ll 1$ is instructive as it leads to

$$\bar{V}^i = V_s^i B/\alpha,$$

showing clearly the necessity of the work of the pumps to stabilize the volume at a finite value in the presence of an osmotic imbalance due to the presence of macromolecules inside the cell (coefficient $B$). The proportionality between the cell volume and the dry mass over a long timescale where synthesis happens is also an important feature [21]. The contribution of the entropy of mixing between the solvent and the biopolymers (coefficient $A$) is quantitatively crucial to set the cell volume. Setting $A$ to zero leads to the unrealistic value $\bar{\phi}_s \simeq 0.23$.

### V. THE OPENING OF GSDMD PORES

A first naive model of the creation of a large number of GSDMD pores could consist in supposing that both permeability parameters $L_p$ an $P$ become very large in (8). With such choice, the parameter $\alpha$ becomes infinitely small since ion pumping is unable to resist to the passive osmotic leaking of ions. In such case, there is therefore no possible steady state and the cell volume swells to infinity. Modulo a mechanical resistance of the cell membrane which is negligible, this would also happen with more refined models [1, 16, 19, 27, 31] which operate with a water flux that follows relation (7). This is not consistent with experimental observations which feature an increase and plateauing of the cell volume for a duration of few minutes upon the creation of GSDMD pores.

Our explanation of this plateau comes from the fact that the water and ion fluxes through the membrane are supported by another contribution compared to (7) which is modified in the following way:

$$\begin{pmatrix} J_w/\rho_w \\ S \\ d\zeta/dt \end{pmatrix} = \begin{bmatrix} L_p + \widehat{L}_p & \sigma\widehat{L}_p & 0 \\ \sigma\widehat{L}_p & P + \sigma^2\widehat{L}_p & \Lambda_{23} \\ 0 & -\Lambda_{23} & \lambda \end{bmatrix} \begin{pmatrix} \Delta p - \Delta\Pi \\ \Delta\mu \\ A \end{pmatrix}. \tag{12}$$

The contributions with hats, containing off-diagonal terms reminiscent of non-selectivity, are related to the presence of the GSDMD pores. The limit where the parameter $\sigma$ is zero corresponds to a selective case since the water and ion fluxes are becoming independent again.

The form (12) is motivated below by considering the pressure drop across a cylindrical hole in a membrane relating the flux of water per unit surface $j_w^p/\rho_w$ to the pressure variation $\Delta p$ in the following way :

$$j_w^p/\rho_w = \mathcal{R}\Delta p,$$

where the effective hydraulic resistance can take two limiting forms depending if the width of the membrane $h$ is much larger than the radius $r$ or not [10]: $\mathcal{R} = r/(3\pi\eta)$ if $r \gg h$ (this is the Sampson limit [28, 29, 32]) and $\mathcal{R} \simeq r^2/(3.75\pi\eta h)$ in the opposite case where $r \ll h$. In the above formulas, $\eta \approx 10^{-3}$Pa.s is the water viscosity. The thickness of the plasma membrane is of the order of $h \approx 5$nm [2] but the typical hydrodynamic radius of GSDMD pores is not clearly determined. It may range from $r \approx 1$nm based on our effective transport measurements (see main text) to $r \approx 20$nm in [24] where only the structure of GSDMD oligomers in the membrane is visualized by AFM, justifying the consideration of the two limiting cases mentioned above. The flux of ions per unit surface $s^p$ across the hole is

$$s^p = \rho_w \phi \mathcal{R} \Delta p,$$

where $\phi$ is the mass fraction of ions in the pore. We neglect the ion diffusive mobility in the pore based on the assumption that an electrochemical constraint controls the mass fraction of ions in the pore sufficiently fast (see Sec. VI for details)

With the purely entropic free energy $f(\phi) = RT(\phi \log\phi - \phi)/M$ describing the ions behavior, the osmotic pressure reads $\Pi = \rho_w(\phi\partial_\phi f - f) = \rho_w RT\phi/M$ and the chemical potential is $\mu = \partial_\phi f = RT/M \log\phi$. We thus obtain:

$$\begin{pmatrix} j_w^p/\rho_w \\ s^p \end{pmatrix} = \mathcal{R} \begin{bmatrix} 1 & \rho_w\phi \\ \rho_w\phi & (\rho_w\phi)^2 \end{bmatrix} \begin{pmatrix} \Delta p - \Delta\Pi \\ \Delta\mu \end{pmatrix},$$

where we made the approximation that variations accross the channel remain small, $\Delta \log\phi \simeq \Delta\phi/\phi$.

In (12), the term $\widehat{L}_p$ can be associated with $S_p\mathcal{R}$ where $S_p = n_p\pi r^2$ is the total surface occupied by the pores over the whole cell membrane and $n_p$ is the total number of pores. The term $\sigma$ can be related to the regulated concentration of ions in a pore $C^* = \rho_w\phi/M$. If $C^*$ is zero, the created pores are selective again as only water can go through them and ions are not driven through the pores by the flow. As soon as $C^*$ increases, there is a certain filtration of the ions through the pores carried by the water flux. The exact value of $C^*$ presumably depends on the pore internal organization and electro-chemical properties [2, 3]. Also, to make the connection between the single channel and the global parameters more quantitative, additional prefactors are expected due to collective hydrodynamic effects between the pores [15].

The exchange model (12) leads to the following modification of (8):

$$\begin{cases} \tau\frac{d(1/\phi_s - 1)}{dt} = (1+n)(A\phi_s^2 + B\phi_s + \chi - 1) - \theta n \log(\chi) \\ \tau\frac{d\chi(1/\phi_s - 1)}{dt} = \theta n(A\phi_s^2 + B\phi_s + \chi - 1) - (n\theta^2 + \epsilon)\log(\chi) - \alpha\epsilon \end{cases} \tag{13}$$

The two additional non-dimensional parameters are

$$n = \frac{\widehat{L}_p}{L_p} \approx \frac{S_p\mathcal{R}}{L_p},$$

representing the membrane permeability due to the GSDMD pores in the membrane compared to that due to the selective aquaporins and

$$\theta = \frac{\sigma}{MC^e} \approx \frac{C^*}{C^e},$$

representing the filtration ratio of the GSDMD pores with respect to the ions which can be expressed as the regulated concentration of ions in the pores compared to the external concentration.

It is difficult to estimate $\theta$ since $C^*$ depends on the molecular details of the pore organization. However, based on an approximate number of pores of $n_p \approx 10^6$ roughly estimated from [24] 10 minutes after the initiation of pyroptosis for a cell of total surface $S_i$, we can estimate $n$ in the two limiting cases where $\mathcal{R}$ represents a large or small pore compared to the membrane thickness. In the first case, we obtain that

$$n \approx \frac{n_p r^3}{3\eta L_p} \approx \frac{10^6 \times 8 \times 10^{-24}}{3 \times 10^{-3} \times 4 \times 10^{-22}} \approx 10^7.$$

The created pores are so large that even a single pore would actually already lead to a similar permeability as all the selective aquaporins. In the second more restrictive case corresponding to our measurements where the pore is considered small compared to the membrane thickness, we have

$$n \approx \frac{n_p r^4}{3.75 h\eta L_p} \approx \frac{10^6 \times 10^{-36}}{3.75 \times 5 \times 10^{-9} \times 10^{-3} \times 4 \times 10^{-22}} \approx 10^2.$$

In this case, an important number of pores ($\approx 10^4$) is necessary to match the permeability due to aquaporins, which is more realistic.

The steady state of (13) found by setting $J_w = S = 0$ corresponds to the system:

$$\begin{matrix} A\bar{\phi}_s^2 + B\bar{\phi}_s + \bar{\chi} + \frac{\alpha\theta n\epsilon}{\theta^2 n + n\epsilon + \epsilon} - 1 = 0 \\ \left(\theta^2 n + n\epsilon + \epsilon\right)\log(\bar{\chi}) + \alpha(n+1)\epsilon = 0. \end{matrix} \tag{14}$$

Solving for $\bar{\chi}$ in (14) we obtain

$$\bar{\chi} = \exp\left(-\frac{\alpha(n+1)\epsilon}{\theta^2 n + n\epsilon + \epsilon}\right), \tag{15}$$

which leads to the second order equation on $\bar{\phi}_s$:

$$A\bar{\phi}_s^2 + B\bar{\phi}_s + \exp\left(-\frac{\alpha(n+1)\epsilon}{\theta^2 n + n\epsilon + \epsilon}\right) + \frac{\alpha\theta n\epsilon}{\theta^2 n + n\epsilon + \epsilon} - 1 = 0. \tag{16}$$

This relation modifies (10) by effectively changing the ion osmotic pressure term that can balance the osmotic pressure due to the presence of macromolecules. The presence of pumping is still necessary since if $\alpha = 0$, we obtain $\bar{\phi}_s = 0$ leading to infinite swelling. However, the effect of the pumping rate is no longer monotone due to a competition between the osmotic and chemical potential contributions to the water flux. Thus a small pumping rate leads to an increase of $\bar{\phi}_s$ while a large pumping rate leads again to infinite swelling. Through both the selective channels and the pores, an increase of ions in the cell will create an influx of water through direct osmosis to dilute the ions. However an increase of the ion internal concentration will also create an efflux of ions to equilibrate the ion concentration across the membrane. Such flux will create a proportional efflux of water through the pores, decreasing the cell volume. The competition between these two phenomena will fix the steady state volume. The cross effect also happens at the level of ions which flux is no longer only proportional to their difference of concentration across the membrane but also contains a contribution proportional to the water flux going through the pores and transporting ions. Both direct osmosis and cross specie transport are proportional to the number of pores, so the full permeabilization of the membrane by the pores is not a trivial limit since both terms are still competing, potentially giving rise to a cell volume plateau.

The solution of (16), which is still a quadratic equation of the same form as (11), can be found explicitly. We consider the limit where $n \gg 1$ which leads to

$$\bar{\phi}_s = \frac{\sqrt{4A\left(1 - \theta\tilde{\alpha} - e^{-\tilde{\alpha}}\right) + B^2} - B}{2A} \quad \text{and } \bar{\chi} = e^{-\tilde{\alpha}} \text{ where } \tilde{\alpha} = \frac{\alpha\epsilon}{\theta^2 + \epsilon}. \tag{17}$$

It is apparent in (17), that both the effective pumping and the effective cell permeability are modified by the presence of the pores. Even in the case where the number of pores is large, there is a range of $0 \leq \theta \leq \theta_m$ where $\theta_m$ only depends on $\alpha$ and $\epsilon$:

$$\alpha\epsilon\theta_m\left(\theta_m^2 + \epsilon\right) = 1 - e^{-\frac{\alpha\epsilon}{\theta_m^2 + \epsilon}}$$

for which $\bar{\phi}_s$ remains finite. We recover the steady state expressions (11) when $\theta = 0$, which corresponds to a full selectivity of the pores since only water can get through in this degenerate case.

### VI. NEGLECTING THE IONS DIFFUSIVE FLUX IN THE GSDMD PORES

We consider that the GSDMD pores can impose a certain concentration of ions along their length. Several mechanisms can contribute to such regulation [2, 12]. Here we do not specify any precise physical mechanism and simply assume that the pore is able to control its internal ion concentration through some electro-chemical process. The evolution of the ion concentration in the pore is then given by:

$$\partial_t C + \partial_x(Cv - D\partial_x C) = \frac{C^* - C}{\tau},$$

where $x \in [0, h]$ is the spatial coordinate along the pore of length $h$, $t$ denotes the time, $C(x,t)$ the concentration of ions in the pore, $v(x,t)$ the velocity of the solvent, $D$ the diffusion coefficient of the ions, $C^*$ is the target concentration that the pore organization selects and $\tau$ is a characteristic time over which this constraint is imposed. At the channel ends, the concentration is imposed: $C(0,t) = C_L$ and $C(h,t) = C_R$. We assume that the velocity of the solvent is given by a Darcy law

$$v = -\frac{\kappa}{\eta}\partial_x p,$$

where $p$ is the hydrostatic pressure and $\kappa$ is the pore permeability. Due to the mass balance in the solvent, $\partial_x v = 0$ and $p = p_L + x(p_R - p_L)/h$ where $p_{L,R}$ are the imposed hydrostatic pressures at the ends of the pore. Thus the velocity of the solvent is constant: $v = -\kappa(p_R - p_L)/(\eta h)$. The steady state concentration then obeys the relation:

$$-l_D^2\partial_{xx}C + l_p\partial_x C + C = C^*,$$

where $l_D = \sqrt{D\tau}$ is a length scale relative to the motion of ions by diffusion and $l_p = v\tau = -\tau\kappa(p_R - p_L)/(\eta h)$ is related to the motion of ions by hydrodynamic transport. Based on the solution of the above linear equation, we then compute the average ion flux through the pore:

$$s = \frac{1}{h}\int_0^h (Cv - D\partial_x C)\mathrm{d}x.$$

Assuming that $l_p \ll l_D$ because $p_R$ and $p_L$ are sufficiently close, we find

$$s = \vartheta^2 \frac{h}{\tau}(C_L - C_R) + \kappa \frac{\vartheta(C_L + C_R - 2C^*)\tanh\left(\frac{1}{2\vartheta}\right) + C^*}{\eta h}(p_L - p_R),$$

where $\vartheta = l_D/h$ is a non-dimensional parameter comparing the diffusive length to the channel length. Further assuming that $\vartheta \ll 1$, the dominating term in the above expression is the hydraulic transport term:

$$s = \frac{\kappa C^*}{\eta h}(p_L - p_R),$$

while diffusion is only contributing to the quadratic order in $\vartheta$. The double limit $|l_p| \ll l_D \ll h$ can always be fulfilled if the reaction fixing the target ion concentration in the pore is sufficiently fast.

---


[1] Ram M Adar and Samuel A Safran. Active volume regulation in adhered cells. *Proceedings of the National Academy of Sciences*, 117(11):5604–5609, 2020.

[2] Bruce Alberts, Rebecca Heald, Alexander Johnson, David Owen Morgan, Martin Raff, Keith Roberts, and Peter Walter. *Molecular Biology of the Cell*. 2022.

[3] Lydéric Bocquet and Elisabeth Charlaix. Nanofluidics, from bulk to interfaces. *Chemical Society Reviews*, 39(3):1073–1095, 2010.

[4] Clotilde Cadart, Larisa Venkova, Pierre Recho, Marco Cosentino Lagomarsino, and Matthieu Piel. The physics of cell-size regulation across timescales. *Nature Physics*, 15(10):993–1004, 2019.

[5] Andrew G Clark, Ortrud Wartlick, Guillaume Salbreux, and Ewa K Paluch. Stresses at the cell surface during animal cell morphogenesis. *Current Biology*, 24(10):R484–R494, 2014.

[6] Torben Clausen. Quantification of na+, k+ pumps and their transport rate in skeletal muscle: functional significance. *Journal of General Physiology*, 142(4):327–345, 2013.

[7] Sybren Ruurds De Groot and Peter Mazur. *Non-equilibrium thermodynamics*. Courier Corporation, 2013.

[8] HY Elmoazzen, JAW Elliott, and LE McGann. The effect of temperature on membrane hydraulic conductivity. *Cryobiology*, 45(1):68–79, 2002.

[9] Paul J Flory. Thermodynamics of high polymer solutions. *The Journal of chemical physics*, 10(1):51–61, 1942.

[10] Simon Gravelle, Laurent Joly, François Detcheverry, Christophe Ybert, Cécile Cottin-Bizonne, and Lydéric Bocquet. Optimizing water permeability through the hourglass shape of aquaporins. *Proceedings of the National Academy of Sciences*, 110(41):16367–16372, 2013.

[11] B. Hille. *Ionic Channels of Excitable Membranes*. Oxford University Press, Incorporated, 1992.

[12] Bertil Hille. Ionic channels in excitable membranes. current problems and biophysical approaches. *Biophysical journal*, 22(2):283–294, 1978.

[13] Else K Hoffmann, Ian H Lambert, and Stine F Pedersen. Physiology of cell volume regulation in vertebrates. *Physiological reviews*, 89(1):193–277, 2009.

[14] Frank C Hoppensteadt and Charles S Peskin. *Modeling and simulation in medicine and the life sciences*, volume 10. Springer Science & Business Media, 2012.

[15] Kaare H Jensen, André XCN Valente, and Howard A Stone. Flow rate through microfilters: Influence of the pore size distribution, hydrodynamic interactions, wall slip, and inertia. *Physics of fluids*, 26(5), 2014.

[16] Hongyuan Jiang and Sean X Sun. Cellular pressure and volume regulation and implications for cell mechanics. *Biophysical journal*, 105(3):609–619, 2013.

[17] Alan R Kay. How cells can control their size by pumping ions. *Frontiers in cell and developmental biology*, 5:41, 2017.

[18] Ora Kedem and Aharon Katchalsky. Thermodynamic analysis of the permeability of biological membranes to non-electrolytes. *Biochimica et biophysica Acta*, 27:229–246, 1958.

[19] JP Keener and James Sneyd. *Mathematical physiology 1: Cellular physiology*. Springer New York, NY, USA, 2009.

[20] Karsten Kruse, Jean-Francois Joanny, Frank Jülicher, Jacques Prost, and Ken Sekimoto. Generic theory of active polar gels: a paradigm for cytoskeletal dynamics. *The European Physical Journal E*, 16:5–16, 2005.

[21] Xili Liu, Seungeun Oh, and Marc W Kirschner. The uniformity and stability of cellular mass density in mammalian cell culture. *Frontiers in Cell and Developmental Biology*, 10:1017499, 2022.

[22] H.F. Lodish. *Molecular Cell Biology*. Cd-Rom. W.H. Freeman, 2000.

[29] Ralph Allen Sampson. Xii. on stokes's current function. *Philosophical Transactions of the Royal Society of London.(A.)*, (182):449–518, 1891.

[30] Jean-Yves Tinevez, Ulrike Schulze, Guillaume Salbreux, Julia Roensch, Jean-François Joanny, and Ewa Paluch. Role of cortical tension in bleb growth. *Proceedings of the National Academy of Sciences*, 106(44):18581–18586, 2009.

[31] Larisa Venkova, Amit Singh Vishen, Sergio Lembo, Nishit Srivastava, Baptiste Duchamp, Artur Ruppel, Alice Williart, Stéphane Vassilopoulos, Alexandre Deslys, Juan Manuel Garcia Arcos, et al. A mechano-osmotic feedback couples cell volume to the rate of cell deformation. *Elife*, 11:e72381, 2022.

[32] Harold L Weissberg. End correction for slow viscous flow through long tubes. *The Physics of Fluids*, 5(9):1033–1036, 1962.

[33] Ewa Zlotek-Zlotkiewicz, Sylvain Monnier, Giovanni Cappello, Mael Le Berre, and Matthieu Piel. Optical volume and mass measurements show that mammalian cells swell during mitosis. *The Journal of cell biology*, 211(4):765, 2015.

### VII. SUPPLEMENTARY FIGURES

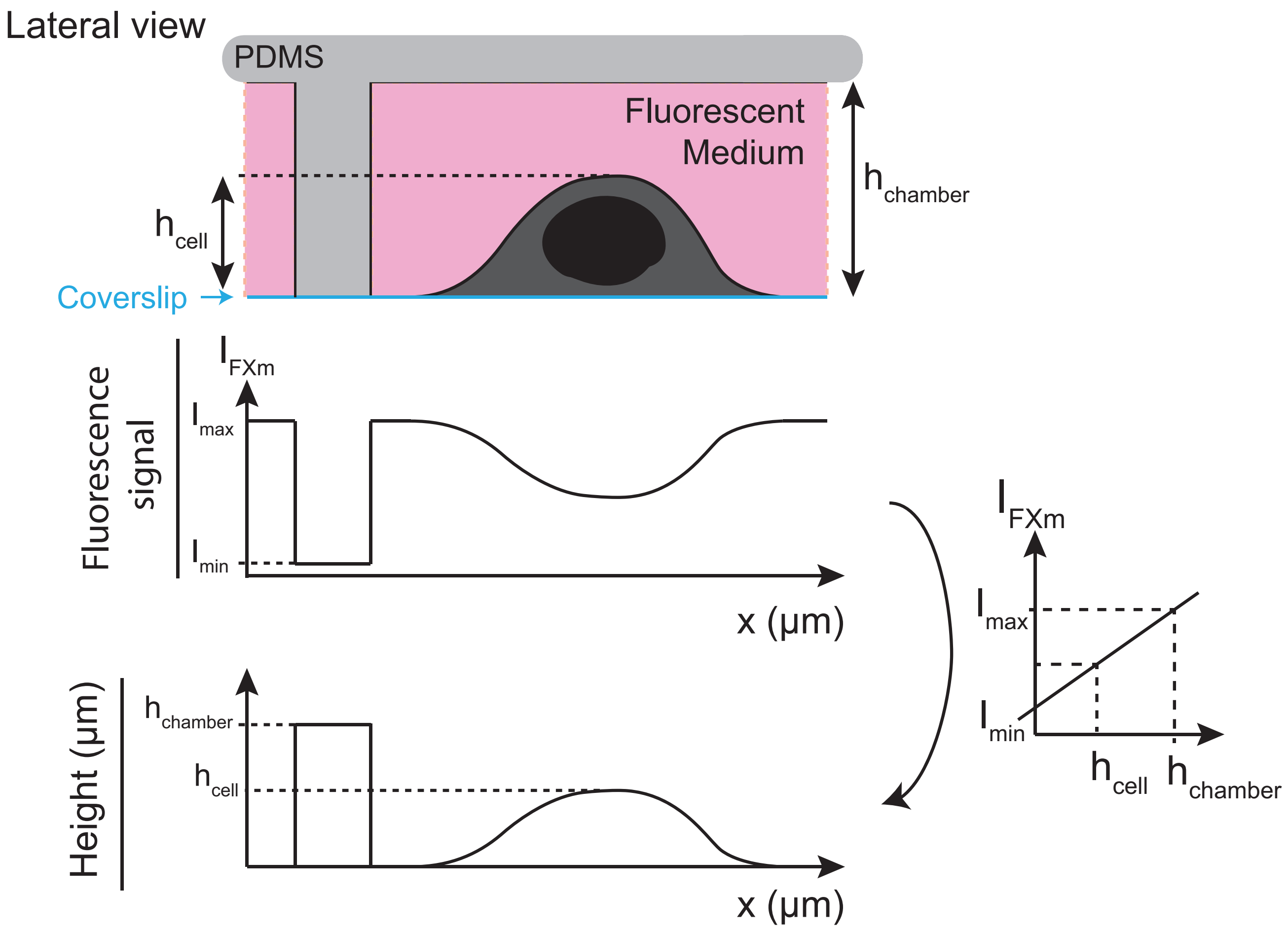


FIG. 1. **Principle of Fluorescence Exclusion microscopy (FXm)**. Lateral view of the setup showing the PDMS chamber in which cells are seeded and surrounded by fluorescent medium. The presence of the cell excludes the dye, creating a local decrease in fluorescence intensity ($I_{\text{cell}}$) compared to the maximal fluorescence ($I_{\text{max}}$) corresponding to the chamber height ($h_{\text{chamber}}$). Epifluorescence microscopy integrates the fluorescence along the whole height of the chamber, the resulting fluorescence intensity profile ($I_{\text{FXm}}$) along the x-axis allows determination of cell height ($h_{\text{cell}}$) relative to chamber height ($h_{\text{chamber}}$). Because fluorescence signal is linearly related to the excluded cell volume height, this provides a quantitative readout of cell volume.

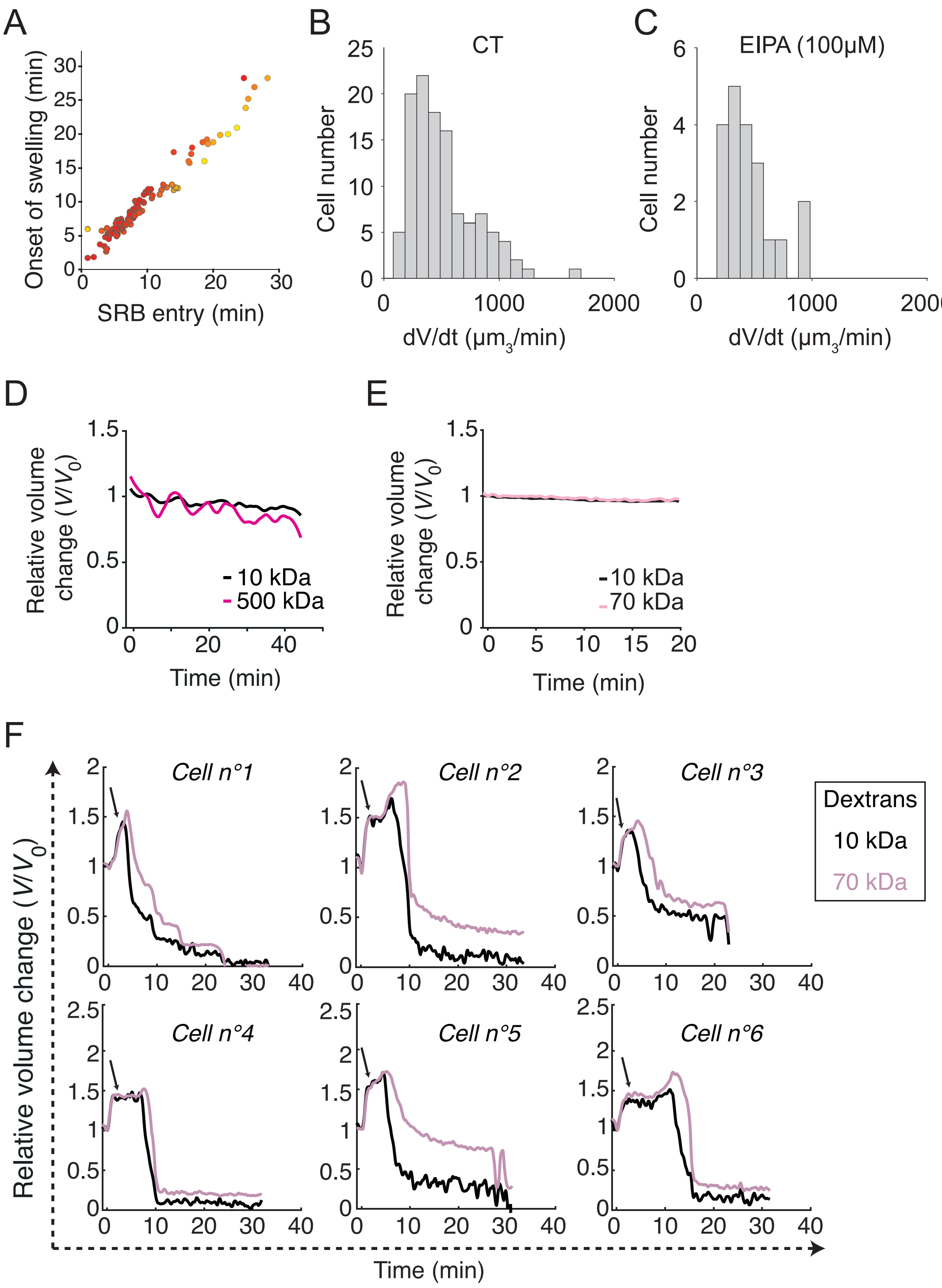


FIG. 2. (A) Correlation between SRB entry and cell swelling onset (pyroptosis phase 1) in individual WT iBMDMs. Color gradient (red to yellow) indicates correlation level (Pearson correlation, $p < 0.001$, $N$=3, $n$=101 cells). (B-C) Distribution of swelling rates ($\Delta V/\Delta t$) during the early swelling phase in untreated (D) and EIPA-treated (E) MEFs. Cells were treated or not with EIPA (500 $\mu$M), an $Na^+/H^+$ exchanger inhibitor, for 1 h. Following treatment and pyroptosis induction via light activation of the optogenetic construct, swelling rates were quantified from the onset of volume increase to the establishment of the steady-state (plateau). (D) Representative normalized volume trajectories of WT iBMDM - without light-activation of pyrotosis - co-incubated with 10 kDa Dextran-AF647 and 500 kDa Dextran-TRITC (relative to the initial volume). (E) Mean normalized volume trajectories of *gsdmd*$^{-/-}$ iBMDMs (relative to the initial volume), co-incubated with 10 kDa Dextran-AF647 and 70 kDa Dextran-FITC. *gsdmd*$^{-/-}$ cells were transfected with the optogenetic construct driving inflammasome activation, light-activated, and their volume was monitored. (F) Example of apparent cell volume dynamics in WT iBMDMs during pyroptosis illustrating the shoulder, consistently observed during swelling (arrow). For (A–B): $N$ = 3, $n$ > 60 cells; for (D): $N$ = 3, $n$ = 114 cells; for (E): $N$ = 1, $n$ = 20 cells; for (G): $N$ = 2, $n$ = 58 cells.

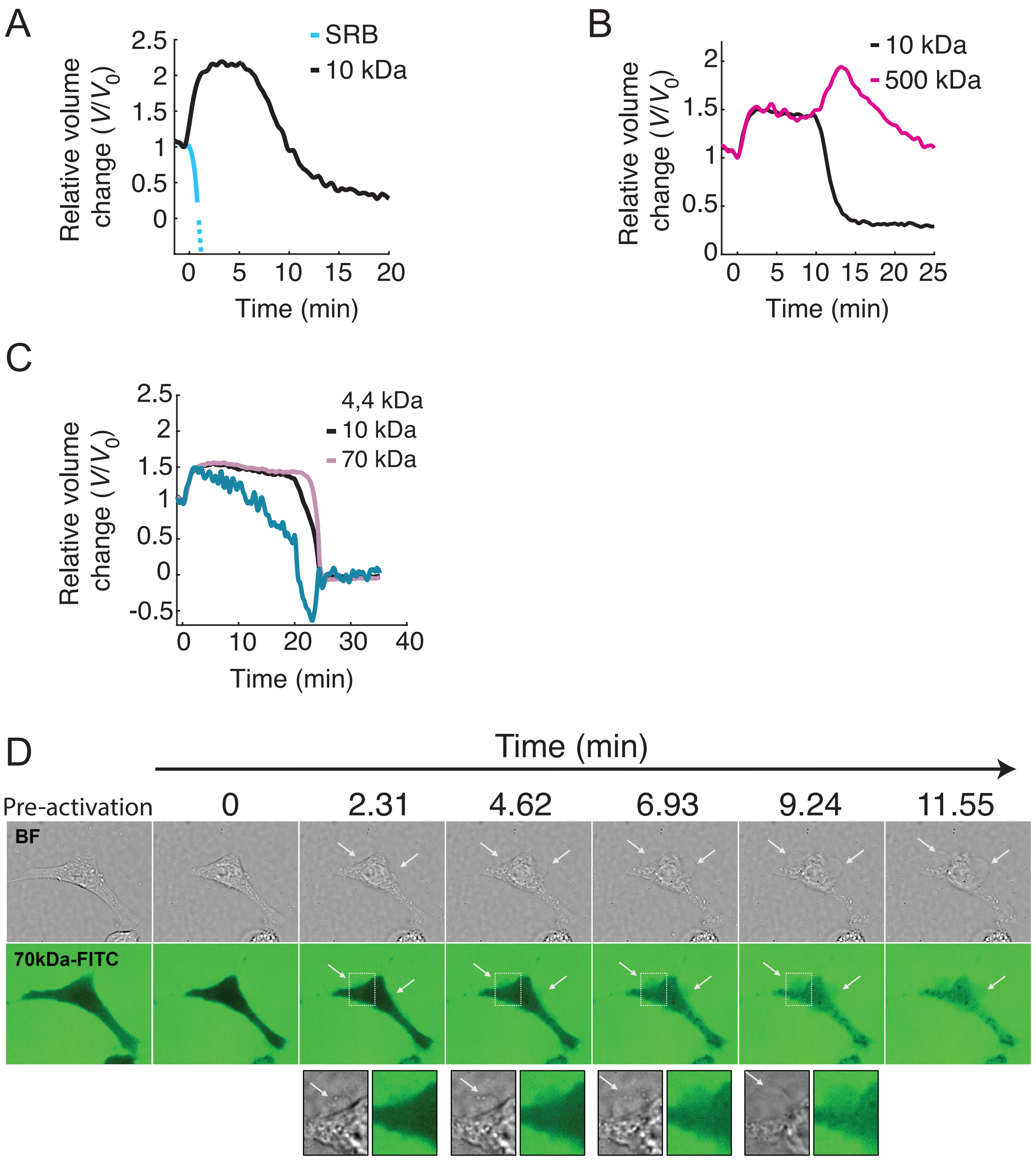


FIG. 3. **Characterization of pyrotosis-driven cell swelling in MEFs.** (A-C) Normalized single-cell apparent volume trajectories (relative to $V_0$ at the onset of cell swelling) of MEFs co-incubated with SRB and 10 kDa Dextran-AF647 (A), with 10 kDa Dextran-AF647 and 500 kDa Dextran-TRITC (B), or with 4.4 kDa Dextran-TRITC, 10 kDa Dextran-AF647, and 70 kDa Dextran-FITC (C). (D) Confocal time-lapse imaging of MEFs undergoing pyroptosis while incubated with 70 kDa Dextran-FITC. Cells were seeded on an ibidi $\mu$-Slide (VI 0.1) and incubated in medium containing 70 kDa Dextran-FITC. Pyroptosis was induced by light activation of the optogenetic construct, and cell morphology along with dextran entry was monitored using a Nikon spinning-disk confocal microscope. Brightfield imaging was used to assess morphology, while FITC fluorescence was detected with a 488 nm laser. Time-lapse acquisition was performed every 15 seconds with a 40× objective. Live-cell imaging was conducted at 37°C with 5% $CO_2$. For (A–B): $N = 3$, $n > 60$, and for C $N$=3, $n$=70 cells.

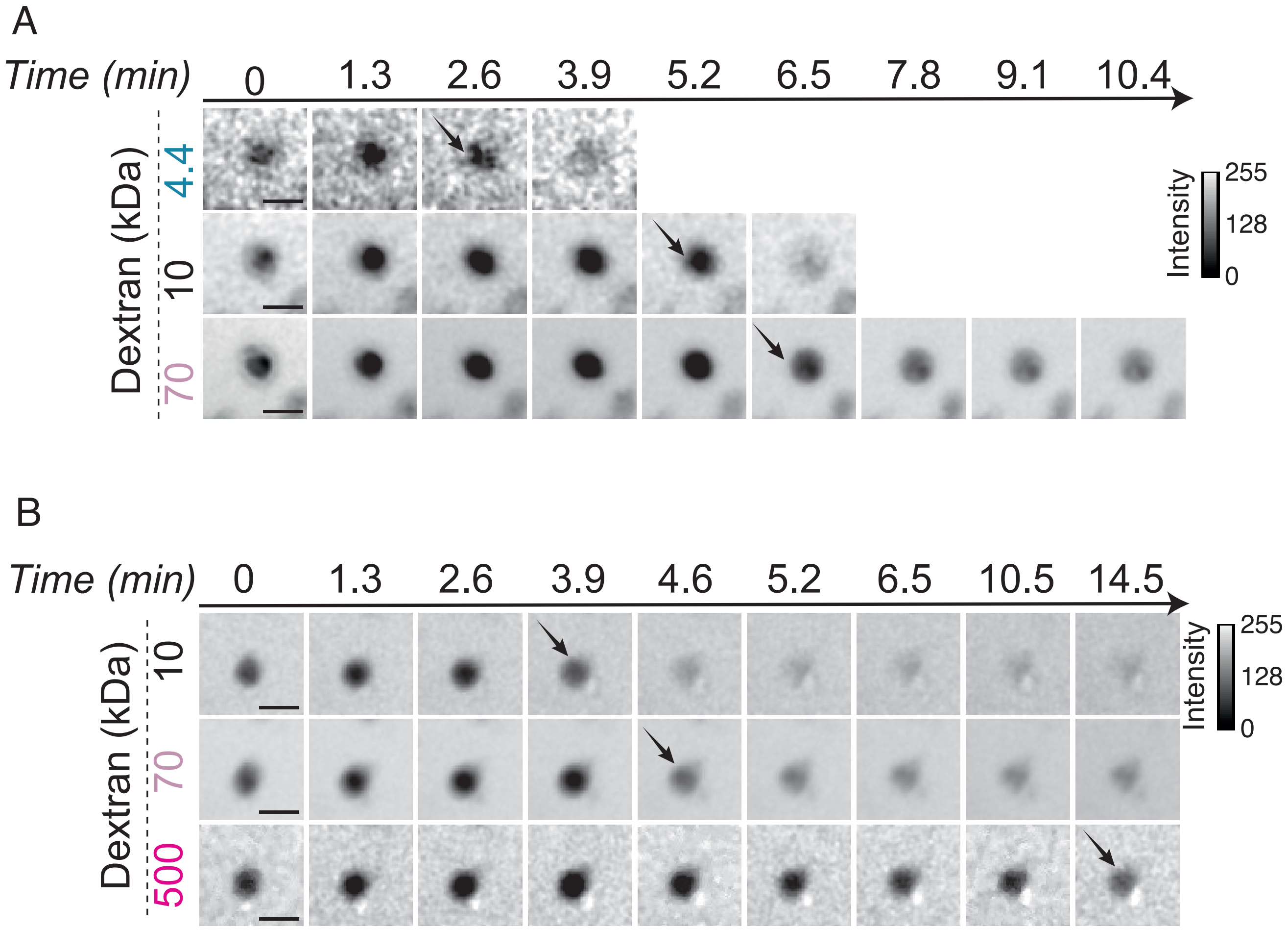


FIG. 4. **Sequential entry of increasing-size Dextrans.** (A,B) FXm time-lapse images of WT iBMDMs undergoing pyroptosis during co-incubation with two sets of fluorescent probes: (A) 4.4 kDa Dextran-TRITC, 10 kDa Dextran-AF647 and 70 kDa Dextran-FITC; (B) 10 kDa Dextran-AF647, 70 kDa Dextran- FITC and 500 kDa Dextran-TRITC. Scale bar, 20 $\mu$m. Black arrows indicate the entry of dyes into the cell. The corresponding normalized single-cell volume trajectories are presented in the main text (Fig.1 E& F). Greyscale: fluorescence intensity

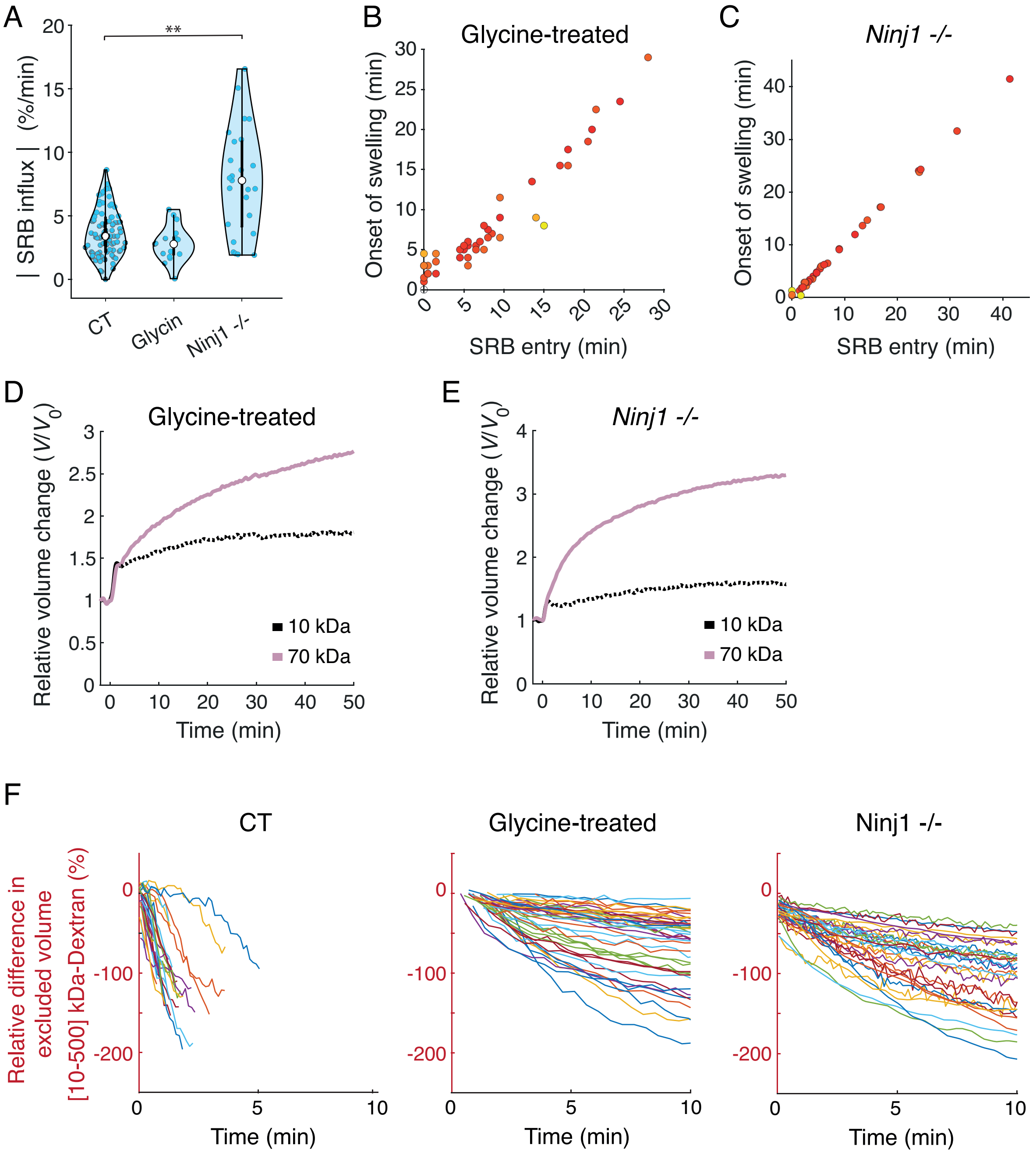


FIG. 5. **Ninj1 and Glycine modulate swelling and dye entry.** (A) Maximal relatifs influx of SRB during the early sweeling phase. See Materials & Methods for parameter computation. Violin plots show single-cell data (colored dots), median (white dot) and SD. Statistical analysis: unpaired t-test ( $^{**}p < 0.01$, $^{***}p < 0.005$, $N = 3$, $n = 33$–$103$ cells across conditions). (B-C) Correlation between SRB entry and swelling onset (pyroptosis phase 1) in WT iBMDMs treated with 20 $\mu$M Glycine (B), or in *Ninj1*$^{-/-}$ iBMDMs (C). Color gradient (red to yellow) indicates correlation level (Pearson correlation, $p < 0.001$, $N = 3$, $n > 60$ cells). (D-E) Representative relative dynamics of apparent cell volume during pyroptosis on WT iBMDMs treated with 20 $\mu M$ Glycine for 2 h (D), and on *Ninj1*$^{-/-}$ iBMDMs (E), monitored by co-incubation with 10 kDa dextran (black) and 70 kDa dextran (pink). (F) Relative difference between excluded volumes measured with 10 kDa and 500 kDa dextran normalized by the volume at onset ($t = 0$). (See Materials & Methods for details). Each curve represents a single cell. Data are shown for WT iBMDMs, Glycine-treated iBMDMs, and *Ninj1*$^{-/-}$ iBMDMs.

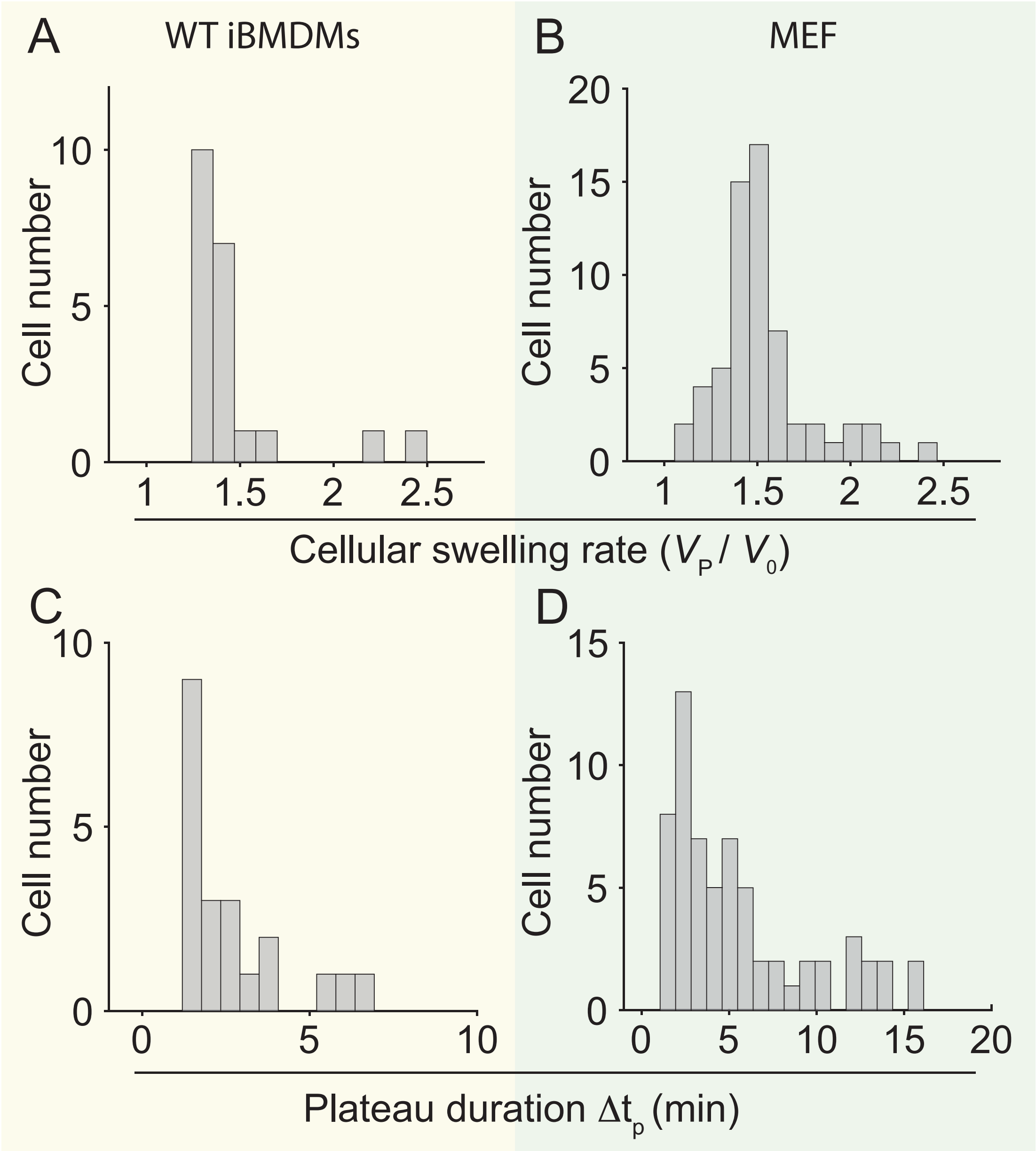


FIG. 6. **Characterization of the volume steady-state.** Distributions of swelling rates required to reach the plateau ($V_{\mathrm{P}}/V_0$) and its duration ($\Delta t_p$) in WT iBMDMs (A–C) and MEFs (B–D). For (A–C): $N = 3$, $n = 21$ cells; for (B–D): $N = 3$, $n = 79$ cells.

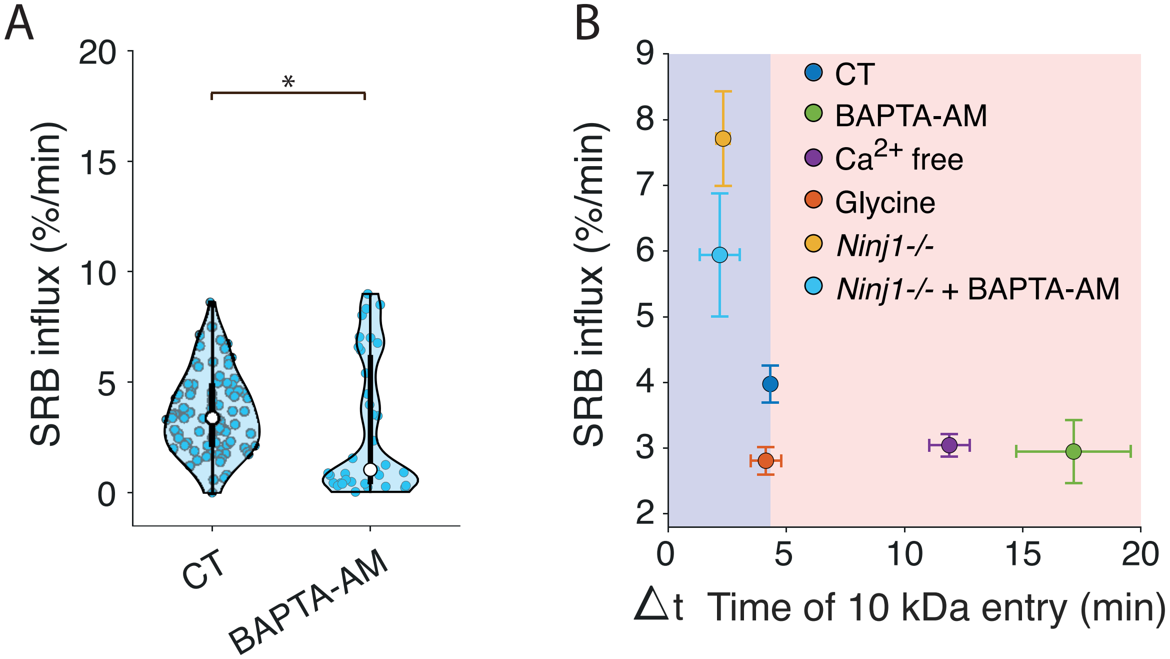


FIG. 7. **Correlation between Sulforhodamine B (SRB) entry and cell volume plateau.** (A) Comparison between untreated (CT) and BAPTA-AM–treated WT iBMDMs of the mean relative SRB influx during the early swelling. Individual cell data are shown (filled dots) with median (white dot) and SD. (B) Mean relative SRB influx as a function of the delay between the onset of swelling ($t_0$) and the onset of 10 kDa Dextran entry, under the indicated conditions on iBMDMs (mean ± SEM). Statistical analysis: Mann–Whitney test, $N = 3$, $n = 87$ CT cells and $n = 31$ BAPTA-AM-treated cells. ($^*$ $p < 0.05$)

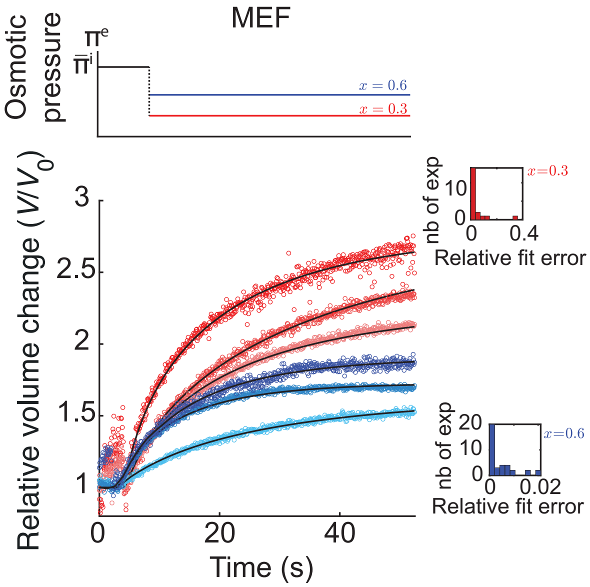


FIG. 8. **Modeling osmotic perturbations in MEFs.** Top: applied osmotic pressure dynamics. Bottom: representative experimental volume trajectories and model fits. The red curves correspond to a $x = 0.3$ dilution factor and the blue $x = 0.6$. The inset plots are bins of the relative squared error between the data and the fit for all the performed experiments in each hypo-osmotic condition ($n = 37$ for $x = 0.6$ and $n = 10$ for $x = 0.3$). See Materials & Methods for fitting parameters.
